# An SE(3)-equivariant Crystal Structure Prediction Framework for Prospective Identification and Development of Bioactive Nucleoside Self-Assembling Materials

**DOI:** 10.64898/2026.05.27.728070

**Authors:** Zheng Wang, Tianyi Wang, Haixian Zhang, Zhenyuan Huang, Chengxi Zhao, Changfu Li, Tiannan Liu, Ding Bai, Xianglong Han, Hang Zhao, Han Wang

**Affiliations:** State Key Laboratory of Oral Diseases & National Center for Stomatology & National Clinical Research Center for Oral Diseases & Research Unit of Oral Carcinogenesis and Management & Chinese Academy of Medical Sciences, West China Hospital of Stomatology, Sichuan University, Chengdu, China; School of Artificial Intelligence, Sichuan University, Chengdu, China; State Key Laboratory of Precision and Intelligent Chemistry, Hefei National Research Center for Physical Sciences at the Microscale, School of Chemistry and Materials Science, University of Science and Technology of China, Hefei, China; Key Laboratory of Green Chemistry & Technology of Ministry of Education, College of Chemistry, Sichuan University, Chengdu, China; State Key Laboratory of Natural and Biomimetic Drugs, School of Pharmaceutical Sciences, Peking University, Beijing, China

## Abstract

Flexible organic small molecules can assemble into supramolecular biomaterials whose properties are intrinsically governed by their crystal structures, yet experimental structure determination remains difficult to scale during molecular modification and materials optimization. Crystal structure prediction (CSP) provides a potential solution, but its prospective power is limited for flexible molecules owing to incomplete conformational sampling and the difficulty of identifying experimentally realized structures from large candidate sets. Here, an SE(3)-equivariant deep-learning workflow, SE3CSP, is developed for organic crystal structure prediction. By learning molecular conformations, unit-cell parameters and packing patterns from experimentally resolved crystal structures and integrating these predictions with MACE-based structure optimization, SE3CSP establishes an end-to-end pipeline from two-dimensional molecular representations to three-dimensional crystal structures. Using nucleosides as representative flexible self-assembling building blocks, SE3CSP achieves an overall prediction accuracy of ∼57%, substantially outperforming the benchmark MACE-based Genarris 3.0 workflow (∼14%). Furthermore, a prospective prediction strategy is developed in which an SE3CSP-predicted density window (± 0.15 g/cm³) is applied prior to energy ranking and structural deduplication, enabling all experimentally realized structure to be consistently ranked within the top 1-2% of all generated candidates. Beyond structure prediction, SE3CSP-derived energy landscapes provide insight into potential single-crystal-to-single-crystal transformations and enable the identification of a nucleoside supramolecular material with dynamic breathing porosity, which is further developed as an adsorptive platform for inflammatory mediator removal with excellent anti-inflammatory performance and biocompatibility. These results establish SE3CSP as a practical framework for prospective CSP and highlight its utility in guiding the discovery and design of bioactive self-assembled materials.

## 1. Introduction

Self-assembled materials formed by flexible organic small molecules are widely used in pharmaceuticals^1,2^, electronics^3-5^ and supramolecular systems^6,7^, and their functions are often closely linked to the crystal structures formed during molecular assembly^8,9^. Experimental crystallographic techniques, including single-crystal X-ray diffraction (SCXRD) and microcrystal electron diffraction (MicroED), provide reliable routes for resolving such structures at high resolution^10^. However, for flexible molecular systems, conformational variability frequently complicates crystallization and limits the scalability of experimental structure determination, particularly during molecular modification and materials screening where repeated characterization is required^11,12^. Consequently, structural evolution during molecular optimization is often difficult to track, limiting the establishment of structure–function relationships and the rational design of self-assembled materials^13,14^. Crystal structure prediction (CSP) provides an alternative route by computationally exploring crystal energy landscapes to identify plausible molecular packings, thereby enabling structural tracking and facilitating structure-guided materials development^15-17^.

Despite rapid advances, CSP continues to face two closely connected challenges: balancing exploration breadth and computational efficiency during conformational sampling for flexible molecules^12,17-20^, and prospectively identifying the experimentally realized form from a large pool of low-energy candidates^8,15,21,22^. A typical CSP workflow comprises conformer search, crystal structure generation, and subsequent optimization and ranking of candidate structures^8,23^. For flexible organic molecules, the first two stages determine whether relevant conformations and packing motifs are sufficiently sampled, whereas the ranking stage governs whether the experimentally relevant structure can be reliably distinguished from numerous energetically competitive alternatives^20^. Current workflows typically rely on density functional theory (DFT) or classical force fields for conformational sampling and lattice-energy evaluation^24-26^. Although DFT-based approaches can accurately describe intermolecular interactions, their computational cost limits scalability, whereas classical force fields tend to favor isolated low-energy conformers that may differ from the conformations stabilized in the crystalline state^27^. Consequently, relevant packing motifs may be absent from the generated candidate pool. More importantly, even when experimentally plausible structures are successfully sampled, lattice-energy ranking alone often fails to reliably identify the correct form because organic crystal energy landscapes commonly contain many structures within a narrow energy window, particularly for flexible and polymorphic systems. Indeed, successive CSP blind tests have repeatedly shown that the experimentally observed structure does not necessarily coincide with the global minimum and may even fall beyond the top 10% of energy-ranked candidates^12,28^. These observations suggest that improving CSP requires not only enhanced conformational and packing sampling, but also more effective prospective ranking strategies incorporating additional physically meaningful constraints beyond lattice energy.

Recent advances in deep learning have begun to address these two challenges by learning structural priors directly from experimentally resolved crystal structures rather than relying solely on exhaustive sampling^29-32^. For the generation problem, OXtal demonstrates that all-atom diffusion models can learn the joint distribution of intramolecular conformations and periodic packing, enabling more experimentally realistic crystal structures to be sampled directly from molecular graphs^33^. For the ranking problem, DeepCSP and MolXtalNet provide an important clue: crystal density can be predicted rapidly and accurately from molecular representations, and can therefore be used to reduce the search space or filter candidate structures before final ranking^23,34^. In particular, MolXtalNet-D achieved a mean absolute error below 2% for density prediction on a large molecular-crystal dataset, supporting density as a robust crystallographic descriptor for CSP filtering^34^. Nevertheless, data-driven generation or density prediction alone does not replace the need for physically grounded structure refinement and energy evaluation^35,36^. These studies instead suggest a more practical route for end-to-end CSP: combining deep-learning-based conformer and packing generation with machine-learned force-field optimization, while using reliably predicted crystallographic descriptors such as density to improve prospective candidate prioritization.

Based on these considerations, an SE(3)-equivariant deep learning workflow, SE3CSP, is developed for organic crystal structure prediction. By learning molecular conformations, unit-cell parameters, and packing patterns from experimentally resolved crystal structures, and integrating these predictions with Message Passing Atomic Cluster Expansion (MACE)-based structure optimization^37^, SE3CSP establishes an end-to-end pipeline from two-dimensional molecular representations to three-dimensional crystal structures (Fig. 1a-c). Using flexible nucleosides, a representative class of self-assembling building blocks, as a testbed, SE3CSP achieves an overall prediction accuracy of ∼57%, substantially outperforming the recently developed representative MACE-based workflow Genarris 3.0 (∼14%)^38^. Furthermore, based on the robust accuracy of deep-learning-based crystal density prediction, a prospective prediction strategy was developed in which an optimized SE3CSP-predicted density window (± 0.15 g/cm³), selected through comparative evaluation, was employed as a pre-filter prior to energy ranking and structural deduplication to improve candidate prioritization. Based on this strategy, all predicted structure closest to the experimental form is consistently ranked within the top 1-2% of all generated candidates. These results demonstrate the potential of SE3CSP for prospective crystal structure prediction and suggest a viable route to guide the design and development of nucleoside-based self-assembling materials, particularly in the context of targeted molecular modification.

**Fig. 1.**
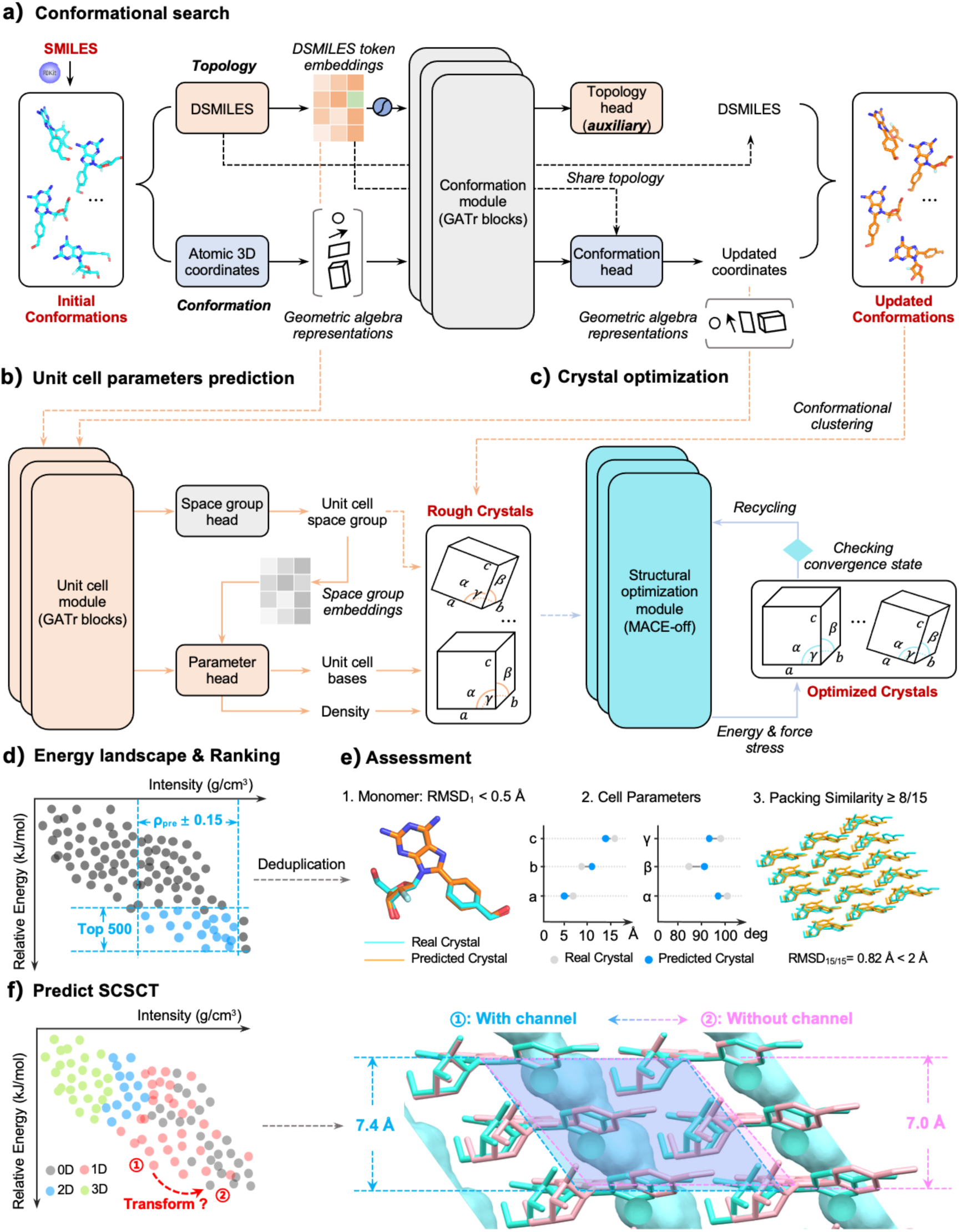
Overview of the SE3CSP workflow and its functionalities. a–c) Schematic illustration of the SE3CSP workflow. Two-dimensional chemical molecular structures are first converted into SMILES representations, followed by initial conformer generation using RDKit. The conformer search module refines molecular conformations, which are then passed to the unit-cell parameter prediction module (a). By learning from organic crystal structures in the Cambridge Structural Database (CSD), this module predicts crystal density, ranges of unit-cell parameters, and likely space groups, enabling the construction of preliminary packed structures (“rough crystals”) (b). These structures are subsequently refined using a batched optimization strategy combined with machine-learned interatomic potentials, with the MACE force field applied to obtain optimized crystal structures (c). d) Energy landscape analysis. A density window defined by the predicted density (ρ_predict_ ± 0.15 g/cm³) was first applied, followed by structural deduplication of the top 500 energy-ranked candidates, to prospectively identify potential crystal structures. e) Evaluation metrics for prediction accuracy, including molecular Root Mean Square Deviation (RMSD), deviations in unit-cell parameters, packing similarity (number of matched molecules), and RMSD_15_. f) The SE3CSP-derived energy landscape accurately predicts a subtle single-crystal transformation, revealing a dynamic breathing porous structure in one nucleoside crystal. RMSD₁ represents the root-mean-square deviation between predicted and experimental molecular conformations, whereas RMSD₁₅ represents the root-mean-square deviation of overlaid 15-molecule packing clusters.

Beyond identifying likely crystal structures, it is noted that SE3CSP-derived energy landscapes can provide insight into possible single-crystal-to-single-crystal transformations. Specifically, when an experimentally observed form is placed on the landscape, lower-energy structures within the SE3CSP-predicted density window (ρ_predict_ ± 0.15 g/cm³) can indicate alternative crystal forms that may be accessible through structural conversion. Using this predictive capability, a candidate nucleoside assembly was identified in which the porous structure exhibits breathing behavior (Fig. 1d-f). This feature was subsequently leveraged to develop an adsorptive material for inflammatory factor removal. Compared with related nucleoside molecules and existing adsorbents, this material shows stronger anti-inflammatory activity, better biocompatibility, and improved performance in both *in vitro* and *in vivo* models. Overall, this work develops a complete workflow for organic crystal structure prediction, enabling structure prediction directly from molecular input and providing a practical framework for the discovery and design of new bioactive self-assembled materials.

## 2. Results

### 2.1. Overview of SE3CSP Complete Workflow

To enable end-to-end prediction of crystal structures from two-dimensional SMILES strings, SE3CSP is constructed as a three-module workflow. The conformational search module (Fig. 1a) generates plausible molecular conformations directly from SMILES representations. The unit-cell parameter module (Fig. 1b) predicts the space group, density, lattice lengths (a, b, c) and lattice angles (α, β, γ), enabling the assembly of preliminary crystal structures. The crystal optimization module subsequently refines both molecular conformations and unit-cell geometry using force-field calculations, yielding optimized crystal structures together with their corresponding energies (Fig. 1c). The first two modules are developed through deep learning on experimentally resolved organic crystal structures from the Cambridge Structural Database (CSD). For the final stage, the MACE force field is employed for crystal structure refinement and energy ranking. Furthermore, motivated by the relatively high accuracy and robustness of deep-learning-derived crystal density prediction, a density-guided prospective ranking strategy was further developed to facilitate prospective crystal structure identification (Fig. 1d). Different from conventional solely energy-based ranking strategies, a density window defined by the deep-learning-predicted crystal density (ρ_predict_ ± 0.15 g/cm³) was first applied to constrain the candidate search space prior to lattice-energy ranking. The remaining structures were subsequently subjected to energy ranking and structural deduplication, thereby enriching experimentally plausible candidates within the generated CSP landscape and facilitating prospective crystal structure identification. To ensure reliable model training and evaluation, a time-split strategy is applied to the CSD dataset to prevent information leakage. Structures that meet the inclusion criteria (Section 4.7.1) and were reported before 2025 are used for training, whereas those from 2025 onward are reserved for validation. In addition, to ensure fair comparison with other methods on the CSD blind test dataset, all related crystal structures are excluded from both sets to avoid cross-contamination.

#### 2.1.1. Predictive Performance in Molecular Conformation and Unit-cell Parameters on the CSD Validation Dataset

To systematically evaluate the SE3CSP’s performance, we developed a comprehensive evaluation framework that assesses the model from both conformational and unit cell parameter perspectives. First, we evaluated the conformation module by calculating the RMSD between the predicted ligand conformations and the corresponding crystal structures in the validation set. For comparison, we also measured the RMSD of the initial conformations generated by RDKit. The overall RMSD distributions for the validation dataset are shown in Fig. **2**a. The average and standard deviation RMSD values for the input and predicted conformations were 1.56 ± 0.92 and 1.36 ± 0.83 respectively, demonstrating that SE3CSP successfully captures subtle conformational variations induced by crystal packing effects and outperforms force-filed based methods in RDKit which predict isolated molecular conformations without accounting for crystal environment constraints.

**Fig. 2.**
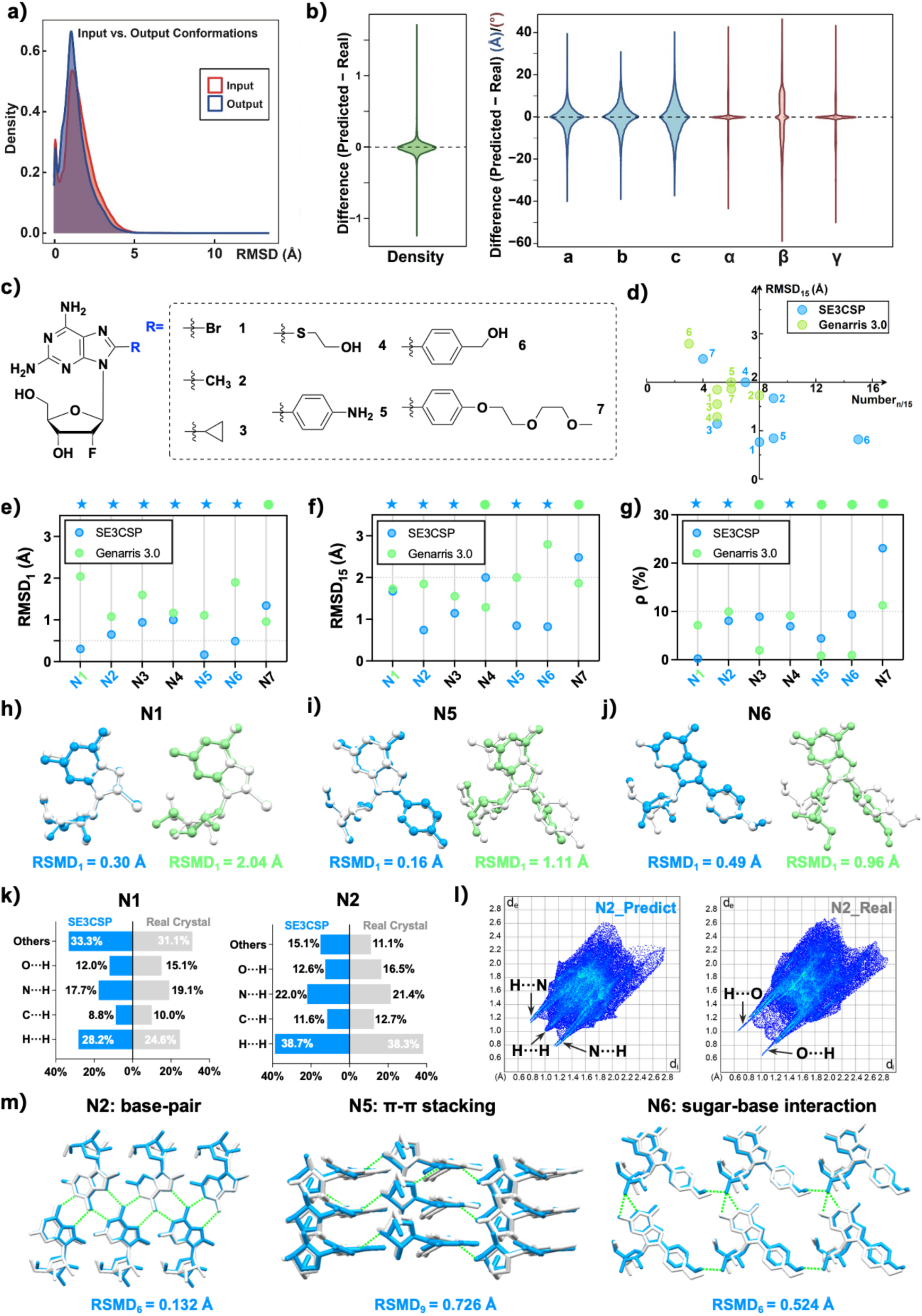
SE3CSP enables rapid and accurate prediction of unit-cell parameters and crystal packing. a) Distribution of RMSD deviations between the updated 2025 organic crystal structures and the corresponding experimental structures before and after SE3CSP prediction. b) Distributions of deviations in crystal density and unit-cell parameters between SE3CSP-predicted structures and experimental structures for the updated 2025 organic crystal set. c) Chemical structures of the candidate 2FA derivatives bearing substitutions at the 8-position, used for evaluation of SE3CSP on flexible nucleoside systems. d) Two-dimensional quadrant plot of packing similarity and RMSD_15_ for structures predicted by SE3CSP and Genarris 3.0. Structures located in the lower-right quadrant represent successful predictions. e–g) Comparison of RMSD_1_, RMSD_15_, and crystal density deviations between SE3CSP- and Genarris 3.0-predicted structures relative to the corresponding experimental structures. h-j) Molecular overlay of predicted and experimental crystal structures for representative cases. Blue stars indicate better performance of SE3CSP, whereas green circles indicate better performance of Genarris 3.0. k) Individual atomic contact percentages obtained from Hirshfeld surface analysis for SE3CSP-predicted and experimental structures. l) Hirshfeld fingerprint plots of SE3CSP-predicted and experimental structures. m) Molecular overlay highlighting key hydrogen-bonding interactions in SE3CSP-predicted and experimental structures. Experimental structures are shown in white, SE3CSP-predicted structures in blue, and Genarris 3.0-predicted structures in green.

We then evaluated the unit cell module using two criteria: the similarity between the normal distributions of the real and predicted lattice parameters (a, b, c, α, β, γ), and the MAE between predicted and actual values. As shown in Fig. **2**, the predicted lattice lengths and lattice angles closely match the distributions of the real data, indicating that SE3CSP has effectively learned the distribution of lattice parameters in natural crystals. The deviation for each lattice parameter was calculated and plotted per crystal, along with the corresponding MAE. Among these, the lattice angle β proved the most difficult to predict accurately. This likely stems from the fact that most natural crystals share the same α and γ values (90°), whereas β exhibits greater diversity—a pattern that has also been observed in a previous study.

#### 2.1.2. Crystal Packing Prediction Accuracy for CSD Blind-Test Candidates

To assess the performance of SE3CSP in predicting crystal packing, twelve compounds satisfying the inclusion criteria were selected from the CSD blind tests (Extended Data Fig. 1a)^12,39-44^. The MACE-based workflow Genarris 3.0 (hereafter Genarris) was included for comparison. A prediction was considered successful when packing similarity analysis identified at least eight matched molecules with an RMSD_15_ below 2 Å. As shown in Extended Data Fig. 1b, SE3CSP correctly predicts 10 out of 12 targets (83%), compared with 6 out of 12 (50%) for Genarris, indicating improved recovery of experimentally observed packing arrangements. On our computing environment (Supplementary Material Section 1.4), the calculations remain computationally tractable: most targets are completed within 18 hours, simpler cases such as Target 6 require approximately 7 hours, and only the most complex system, Target 12, approaches 24 hours (Supplementary Table 1). Relative to DFT-based CSP protocols reported in blind tests, this corresponds to a reduction in computational cost by approximately four to five orders of magnitude. Although the RMSD_1_ and RMSD_15_ values are higher than those typically obtained from DFT-based approaches, this does not prevent identification of the correct packing in most cases. For successful predictions, the best-matched structures generally exhibit RMSD_1_ values below 0.5 Å, with deviations in unit-cell parameters within 10%, indicating good agreement with experiment at the level relevant for structure assignment (Extended Data Fig. 1c-d). Further comparison shows that SE3CSP improves the prediction of RMSD_1_, crystal density (ρ), and the more variable unit-cell parameter β relative to Genarris (Extended Data Fig. 1e-f and Supplementary Table 2). In particular, density prediction is consistently more accurate across all systems considered. This is relevant for self-assembled materials, where features such as porosity are closely linked to crystal density, and therefore benefit from improved density prediction. Overall, SE3CSP provides improved recovery of crystal packing compared with an existing MACE-based workflow, while maintaining a computational cost compatible with routine CSP calculations.

### 2.2. SE3CSP-guided Development of New Nucleoside Supramolecular Materials

#### 2.2.1. Predictive Performance for Flexible Nucleoside Crystals

SE3CSP was next evaluated on chemically modified analogues to determine whether crystal structures derived from site-specific substitution of a common scaffold could be accurately predicted. Flexible nucleosides were selected as a representative test system because they combine strong self-assembly propensity with substantial conformational flexibility. Our previous work shows that the nucleoside 2FA exhibits strong self-assembly capability, and structural variations could be induced through substitution at the 8-position, leading to diverse assembled structures^45-47^. This system therefore provides a suitable platform to assess the ability of SE3CSP to capture structural changes arising from chemical modification. Accordingly, seven 2FA derivatives bearing substituents of increasing size and flexibility at the 8-position were then selected as test targets (Fig. 2c and Supplementary Fig. 2-5). As shown in Fig. 2d-g, SE3CSP correctly recovers 4 out of 7 targets (57%), whereas Genarris recovers 1 out of 7 (14%). These results indicate improved performance in predicting crystal structures of modified nucleosides. At the molecular level, the predicted conformations are largely consistent with experiment. Minor deviations are observed in sugar puckering and substituent orientation at the 8-position, while key torsional relationships within the nucleoside framework are preserved (Fig. 2h and Extended Data Fig. 2a-d). At the packing level, Hirshfeld surface analysis shows that the relative contributions of intermolecular contacts are well reproduced (Fig. 2k and Extended Data Fig. 2e). The corresponding fingerprint plots display similar overall features, with only limited differences in the sharpness of characteristic spikes. The dominant interaction motifs in nucleoside crystals—including base pairing, π–π stacking and sugar–base interactions—are consistently recovered (Fig. 2l and Supplementary Fig. 6).

Notably, SE3CSP provides consistently accurate predictions of crystal density for both CSD blind-test candidates and nucleoside systems, consistent with previous observations that density is generally one of the most reliably predicted crystal parameters in CSP workflows (Table 1). For correctly predicted structures, deviations between predicted and experimental densities remained within 10%. Consistent with previous deep-learning-based CSP studies, including DeepCSP and MolXtalNet, which identified crystal density as one of the most robustly predictable crystallographic descriptors, these results further support the use of density as a physically meaningful constraint for candidate prioritization. Given that the densities of most organic molecular crystals typically fall within the range of ∼1–3 g/cm³, this level of prediction error corresponds to an uncertainty on the order of 0.1–0.3 g/cm³. Guided by this range, different density tolerance windows were comparatively evaluated, and a threshold of 0.15 g/cm³ was found to provide the optimal balance between candidate retention and prospective identification performance for subsequent structure screening (Supplementary Table 3). With this additional constraint, predicted structures within the density window were further prioritized through energy ranking and structural deduplication. Using this strategy, 7 of the 10 correctly predicted CSD blind-test candidates were successfully prioritized within the top 1% of all generated structures (top 25 of 2,500 candidates). Similarly, for nucleoside systems, 3 of the 4 correctly predicted structures were also ranked within the top 2% of generated candidates (Table 2). These results suggest that density-guided filtering can substantially improve the prospective identification of experimentally relevant crystal structures from broad CSP landscapes. Importantly, SE3CSP therefore does not simply enhance structural sampling, but also strengthens prospective structure identification, which is particularly valuable for flexible molecular systems where numerous competing low-energy packings can otherwise complicate prospective CSP.

**Table 1.** Experimental crystal density, SE3CSP-predicted density, and the corresponding deviation (pred. − expt.) (g/cm³).

| Molecule | Experimental $\rho$ | SE3CSP Predicted $\rho$ | Deviation |
| --- | --- | --- | --- |
| Target 1 | 1.338 | 1.263 | 0.075 |
| Target 2 | 1.606 | 1.750 | -0.144 |
| Target 3 | 1.484 | 1.423 | 0.061 |
| Target 4 | 1.528 | 1.529 | -0.001 |
| Target 5 | 1.669 | 1.608 | 0.061 |
| Target 6 | 1.152 | 1.165 | -0.013 |
| Target 7 | 2.528 | 2.564 | -0.036 |
| Target 8 | 1.479 | 1.471 | 0.008 |
| Target 9 | 1.385 | 1.356 | 0.029 |
| Target 10 | 1.837 | 1.826 | 0.011 |
| Target 11 | 1.566 | 1.682 | -0.116 |
| Target 12 | 1.478 | 1.525 | -0.047 |
| N1 | 1.725 | 1.732 | -0.007 |
| N2 | 1.464 | 1.388 | 0.076 |
| N3 | 1.450 | 1.413 | 0.037 |
| N4 | 1.493 | 1.487 | 0.006 |
| N5 | 1.475 | 1.431 | 0.044 |
| N6 | 1.375 | 1.396 | -0.021 |
| N7 | 1.290 | 1.431 | -0.141 |

**Table 2.**
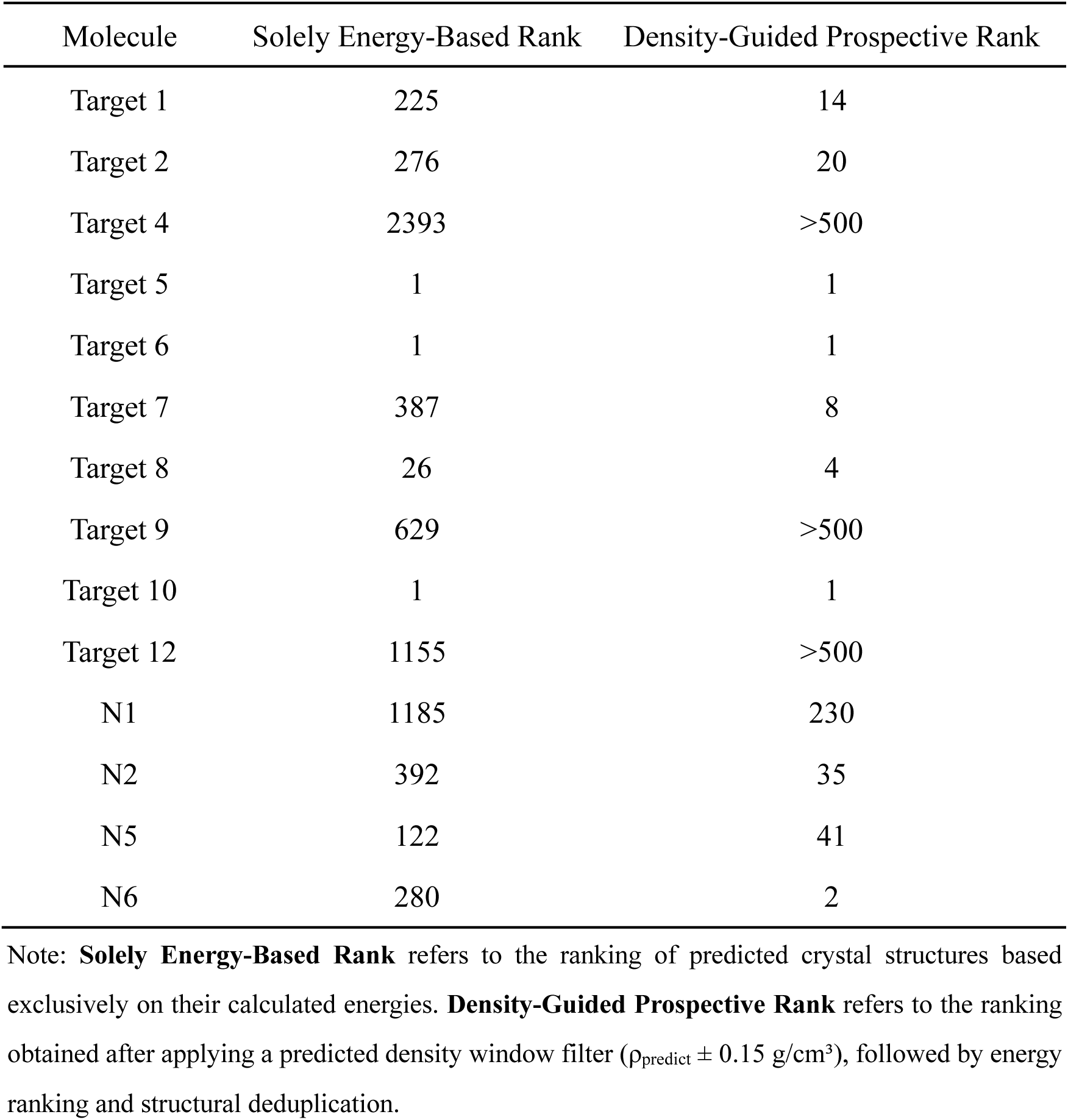
Rank and RMSD₁₅ of the predicted structure with the highest packing similarity to the experimental structure among the top 500 candidates generated by solely energy-based and density-guided prospective ranking strategies.

Overall, SE3CSP provides faster and more accurate prediction of nucleoside crystal structures than conventional workflows such as Genarris, and shows promise for the prospective prediction of flexible self-assembling molecular crystals. This capability may facilitate routine evaluation of how chemical modification alters self-assembled structures under routine laboratory conditions.

#### 2.2.2. SE3CSP-enabled Discovery of a New Nucleoside Self-Assembling Material 8BA with Dynamic Breathing Porosity

Having established the prospective prediction capability of SE3CSP for organic crystal structures, its potential to predict polymorphism and single-crystal transformations was further investigated. In this context, the crystal energy landscape provides a useful representation of the relative stability of alternative packing arrangements and allows identification of low-energy structures that may be accessible from a given crystalline state.

Among the seven nucleoside derivatives examined above, the system with the highest prediction accuracy, 8BA (denoted N6), was selected for detailed analysis. At the molecular level, the 8BA molecule adopts a *syn* conformation between the nucleobase and sugar moiety, stabilized by intramolecular hydrogen bonding (O5’– H···N3, C2’–H···O5’ and C2’–H···N3), with a *C2’-endo* sugar pucker (Fig. 3a and Supplementary Table 4). SE3CSP reproduces this conformation with high accuracy, with only minor deviations in the orientations of the 5’-OH and 16-OH groups, and shows clear improvement over Genarris (Fig. 2j). At the packing level, SE3CSP achieves full recovery of the experimental structure with 15/15 molecules matched and an RMSD_15_ of 0.82 Å, whereas the best Genarris prediction matches only 3 molecules with an RMSD_15_ of 2.794 Å. Powder X-ray Diffraction Patterns (PXRD) further confirm that the SE3CSP-predicted structure is in closer agreement with experiment (Fig. 3b).

**Fig. 3.**
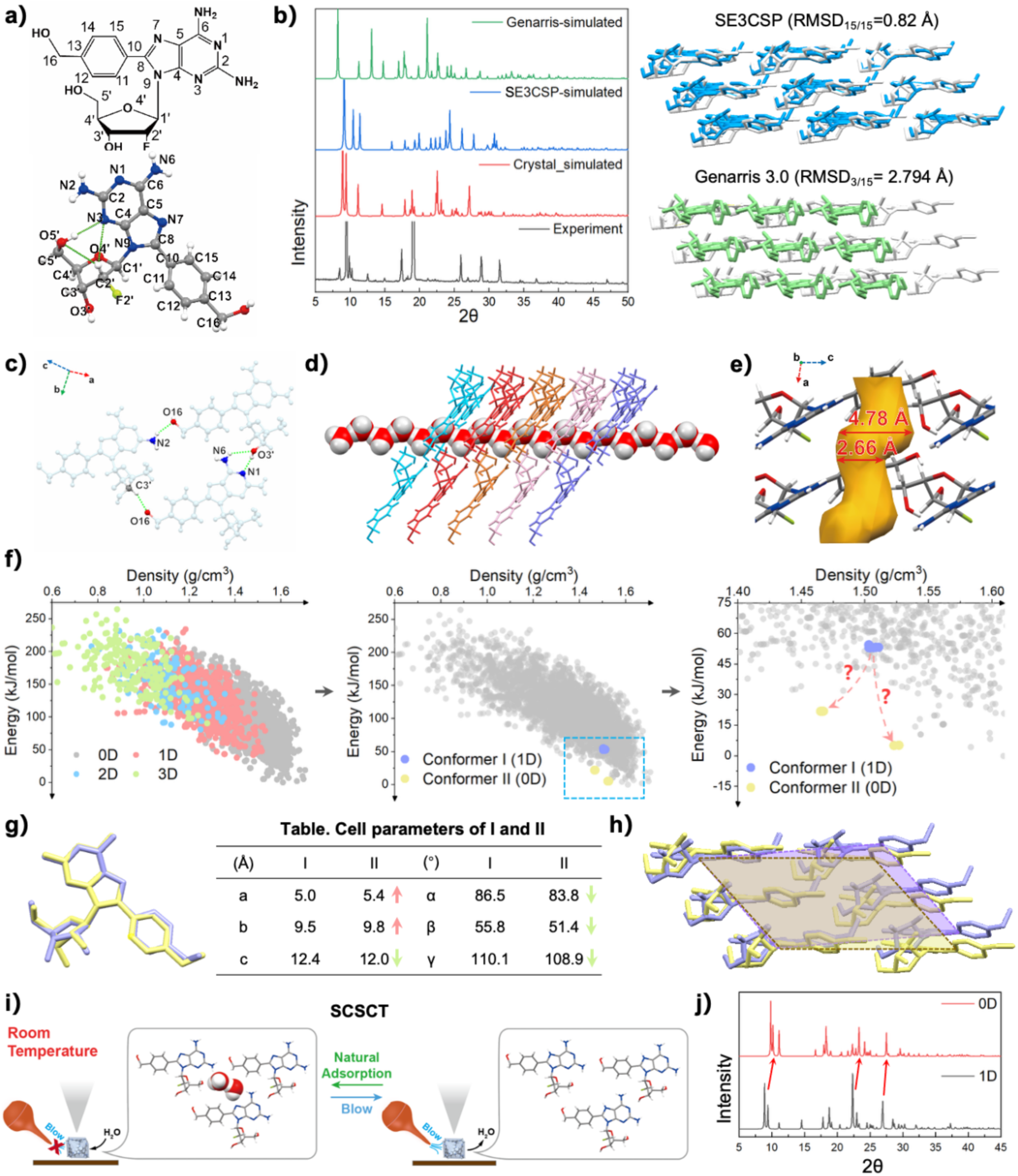
SE3CSP enables identification of dynamic breathing porosity in nucleoside 8BA. a) Chemical structure of 8BA and its experimentally determined crystal structure. b) PXRD patterns of 8BA: experimental PXRD, PXRD simulated from the single-crystal structure, and PXRD patterns of structures predicted by SE3CSP and Genarris 3.0 (left); molecular overlays of predicted and experimental structures (right). Experimental structures are shown in white, SE3CSP predictions in blue, and Genarris 3.0 predictions in green. c–e) Molecules assemble into trimeric units through intermolecular hydrogen bonding, forming water-channel-like structures (c). Linear arrangement of water molecules within the 1D channel (d). Channel structure highlighted in orange (e). Atoms are colored as follows in Fig. 3e: red, oxygen; blue, nitrogen; gray, carbon; white, hydrogen; green, fluorine. f) SE3CSP-derived energy landscape of 8BA annotated using ZOE++. The best-predicted structure (state I, purple) exhibits a 1D porous channel consistent with the experimental structure, while a lower-energy structure (state II, yellow) without porosity is identified. The relative positions of the two states suggest a possible structural transition. g) Molecular overlay and comparison of unit-cell parameters between state I and state II. h) Structural illustration obtained by overlaying state II onto state I, highlighting the collapse of the 1D channel. i) Experimental validation of single-crystal transformation in 8BA, showing reversible insertion and removal of guest water molecules within the 1D channel. j) PXRD patterns simulated from the single-crystal structures of 8BA in the left (1D) and right (0D) panels of i.

The experimental crystal structure of 8BA adopts the P1 space group and forms a porous framework through intermolecular hydrogen bonding (N2–H2A···O16, C3’– H···O16, O3’–H···N1 and N6–H6A···O3’), in which molecules assemble into trimeric units (Fig. 3c). These units stack via π–π interactions to form one-dimensional channels along the **a** axis, with water molecules confined within the channels (Fig. 3d-e). This channel-like architecture resembles previously reported nucleoside systems, although the latter form similar structures through tetrameric rather than trimeric assemblies, highlighting structural diversity in nucleoside-based self-assembly (Supplementary Fig. 7).

In CSP, the crystal energy landscape describes the ensemble of possible crystal packings and their relative thermodynamic stabilities, and is widely used to analyze polymorphism, structural transitions and structure–property relationships. By projecting experimentally relevant structures onto this landscape, low-energy competing states potentially accessible through structural transformation can also be identified. Furthermore, annotation of pore topology and dimensionality (0D, 1D, 2D or 3D) using tools such as ZEO++ enables the correlation of structural transitions with changes in accessible porosity, providing additional insight into breathing behavior and pore evolution across the energy landscape. Based on this, projection of the predicted structure onto the energy landscape shows that the best-matched structure (state I) corresponds to a porous phase consistent with experiment. Within the SE3CSP-predicted density window (ρ_predict_ ± 0.15 g/cm^3^), a non-porous lower-energy structure (state II) is identified (Fig. 3f). This structure retains the P1 symmetry and a similar molecular conformation, but lacks the porous channel. Analysis of the unit-cell parameters indicates that this difference arises from a decrease in the **c** axis and lattice angles (α, β and γ), accompanied by an increase in **a** and **b**, resulting in a more inclined unit cell and collapse of the pore (Fig. 3g-h). This prediction suggests that the water molecules within the channels are not rigidly confined and that a lower-energy collapsed phase may be accessible upon water removal. Subsequent single-crystal transformation experiments confirm this hypothesis. Upon gentle gas flow at room temperature, a structural transformation to a state closely resembling state II is observed, while re-exposure to ambient conditions restores the original porous structure (state I) within approximately 20 minutes (Fig. 3i). Powder diffraction patterns show a systematic shift of characteristic peaks to higher 2θ values during the transition from state I to state II, consistent with the contraction of intermolecular distances and the change in unit-cell geometry (Fig. 3j). These results indicate that guest water molecules are essential for stabilizing the porous framework of 8BA and that the channels remain dynamically accessible, allowing reversible guest exchange without loss of crystallinity. Such behavior, characteristic of soft porous crystals, suggests that the channels may provide accessible interfaces for interaction and potential adsorption of hydrophilic species in aqueous environments.

These results demonstrate that analysis of SE3CSP-derived energy landscapes can facilitate identification of single-crystal transformations and provide insight into dynamic structural behavior. In particular, the reversible breathing behavior observed in 8BA highlights the potential of this approach for predicting responsive structural changes in self-assembled molecular crystals and suggests an expanded role for energy landscapes in CSP beyond static structure prediction.

#### 2.2.3. Efficient Adsorption of Inflammatory Factors Facilitated by Breathing Porosity in 8BA

Dynamic porous behavior can facilitate guest uptake in soft porous crystals, suggesting that such structures may be advantageous for adsorption-based removal of inflammatory mediators, which is increasingly recognized as a useful anti-inflammatory strategy^48-50^. The precursor nucleoside scaffold of 8BA was previously shown to possess anti-inflammatory activity, and the nucleoside-based structure of 8BA further suggested possible interactions with nucleic-acid-related inflammatory factors such as cfDNA (cell-free DNA). The breathing porous structure identified in 8BA therefore prompted further examination of its adsorption behavior towards inflammatory factors.

Optical microscopy and scanning electron microscopy (SEM) revealed that 8BA self-assembled crystals exhibited distinct blocky and layered structures (Fig. 4a). Under high-angle annular dark-field scanning transmission electron microscopy (HAADF-STEM), these structures displayed porous layered characteristics (Fig. 4b). Zeta potential measurements demonstrated that 8BA crystals exhibited weak positive charge under both neutral (pH 7.4) and acidic (pH 5.0) conditions, with overall zeta potential values lower than those of the classical cationic adsorbent PAMAM-G3 (polyamidoamine generation 3). In contrast, 2FA and non-porous 2-Amino-8-phenyl-2’-deoxy-2’-fluoro-D-adenosine (8PA) crystals exhibited weak negative charge at both pH conditions (Supplementary Fig. 8-9). The binding capacities of chlorhexidine, a clinically used maintenance drug for periodontitis, PAMAM-G3, 2FA, 8PA and 8BA toward cfDNA, TNF-α, LPS and IL-6 were evaluated at concentrations of 2 mM and 1 mM. The 2 mM materials demonstrated slightly higher binding capacities compared to 1 mM. Notably, 8BA exhibited the highest binding capacity, reaching up to 7-fold that of 2FA or 8PA, and marginally surpassing the binding capacity of PAMAM-G3. The binding capacity of 8BA toward TNF-α was significantly higher than that of the PAMAM-G3 group (Fig. 4c and Supplementary Fig. 10-17). Then, RAW264.7 cells were stimulated with LPS or cfDNA, simultaneously treated with 1 mM or 2 mM 8BA, and cultured for 24 hours. Cell supernatant was collected, and TNF-α and IL-6 levels were measured using ELISA (enzyme-linked immunosorbent assay) kits. Under baseline conditions without inflammatory stimulus, TNF-α and IL-6 levels in the supernatant remained low. Following LPS or cfDNA stimulation, TNF-α and IL-6 levels increased 3-4 fold. Administration of 8BA significantly reduced TNF-α levels, while IL-6 levels decreased to less than 50% of the stimulated group (Fig. 4d-e).

**Fig. 4.**
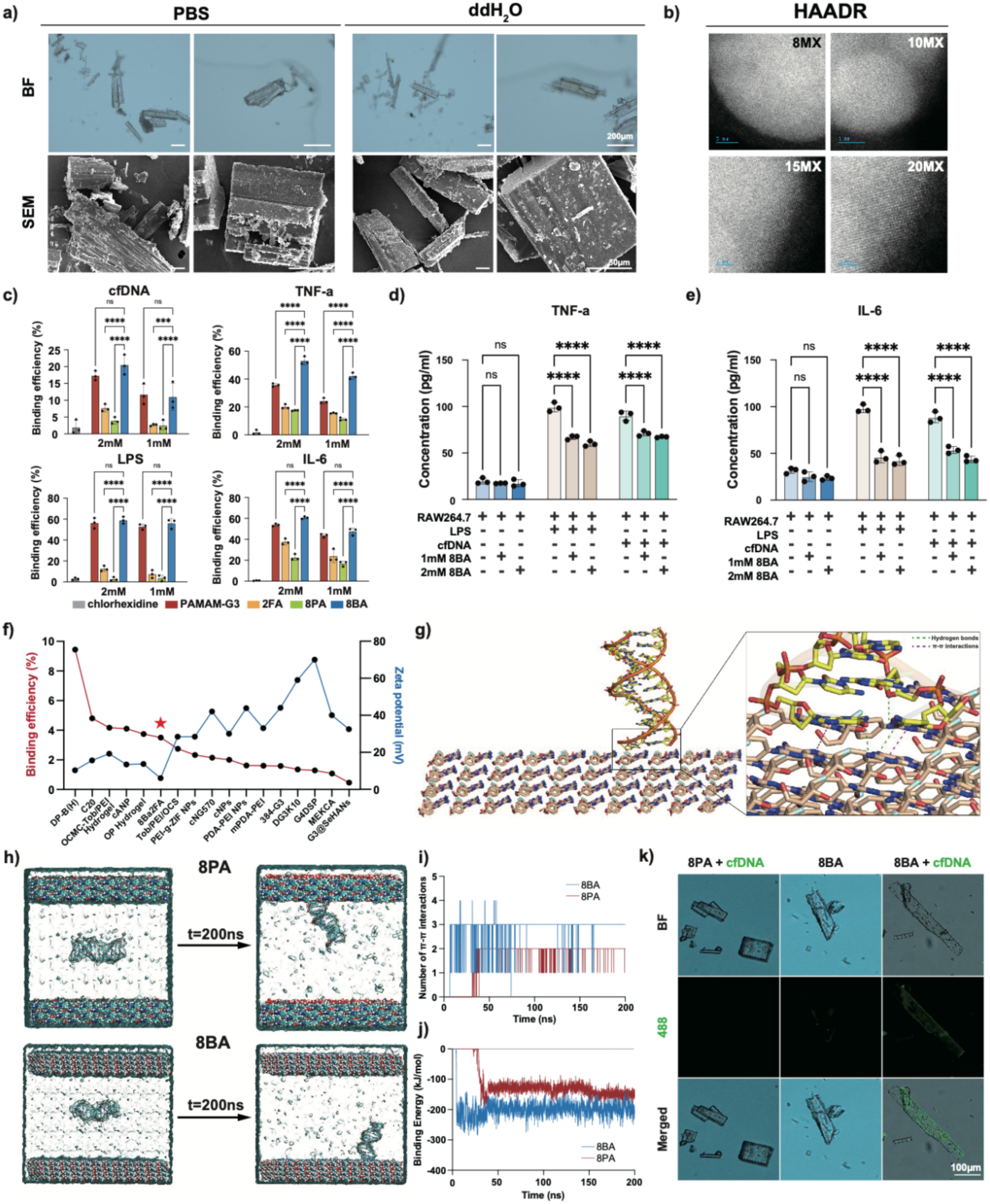
Breathing porous structure supports efficient adsorption of inflammatory factors by 8BA. a) Representative optical microscopy images (upper panel, scale bar = 200 μm) and scanning electron microscopy (SEM) images (lower panel, scale bar = 50 μm) of 8BA materials dissolved in PBS (left) or ddH₂O (right). b) Representative HAADR images of 8BA materials dissolved in PBS. c) Bar plot showing adsorption efficiency of chlorhexidine, PAMAM-G3, 2FA, 8PA and 8BA toward cfDNA, TNF-α, LPS and IL-6 at concentrations of 2 mM or 1 mM. d) Bar plot showing TNF-α level in RAW264.7 cell supernatant after treatment with LPS, cfDNA, 1 mM 8BA or 2 mM 8BA, n = 3. e) Bar plot showing IL-6 level in RAW264.7 cell supernatant after treatment with LPS, cfDNA, 1 mM 8BA or 2 mM 8BA, n = 3. f) Line plot showing correlation between cfDNA adsorption efficiency and zeta potential for various materials, with 8BA marked with asterisk. g) Molecular dynamics simulation of cfDNA binding on 8BA material surface. Green dashed lines indicate hydrogen bonds, red dashed lines indicate π-π interactions. h) Structural changes of 8PA and 8BA systems before and after molecular dynamics simulation. i) Quantification of π-π interactions in 8BA (blue) and 8PA (red) systems. j) Binding energy comparison between 8BA (blue) and 8PA (red) systems. k) Representative microscopy and fluorescence images of cfDNA (green) adsorption by 8PA and 8BA, scale bar = 100 μm. Statistical analyses were performed using two-way ANOVA with Tukey’s multiple comparisons test in (b, c, d and e). * p < 0.05, ** p < 0.01, *** p < 0.001, **** p < 0.0001. In (b, c, d and e), bar graphs are shown as mean ± SD.

A literature review was conducted to analyze the correlation between cfDNA binding capacity and zeta potential across different binding materials (zeta > 0)^51-66^. Among all materials analyzed, 8BA exhibited the lowest zeta potential while ranking among the top 6 in cfDNA binding capacity (Fig. 4f). Molecular dynamics simulations were performed to investigate the binding processes of porous 8BA and non-porous 8PA toward cfDNA. Significant binding of cfDNA molecules was observed on the porous surfaces of 8BA (Fig. 4g and Supplementary Fig. 18). The hydrophobic interactions between cfDNA and material molecules in both 8BA and 8PA systems stabilized after approximately 100 ns, with average values of 1.51 ± 0.60 and 1.22 ± 0.47, respectively (Fig. 4h and Supplementary Fig. 19a). The π-π interactions between cfDNA and the 8BA surface were relatively stronger compared to the 8PA system, with average values of 2.98 ± 0.19 and 1.94 ± 0.23, respectively (Fig. 4i). The hydrogen bond counts between cfDNA and material molecules in both systems were relatively weak, with average values of 0.84 ± 0.76 for 8BA and 0.31 ± 0.55 for 8PA (Supplementary Fig. 19b). The binding energy between cfDNA and material molecules showed distinct kinetic profiles in the two systems. In the 8BA system, the binding energy rapidly increased within 10 ns, indicating fast and stable binding of cfDNA onto the material surface, with an average binding energy of -197.89 ± 18.07 kJ/mol. In contrast, the 8PA system exhibited a delayed increase in binding energy at approximately 40 ns, with an average value of -141.93 ± 15.90 kJ/mol (Fig. 4j). Overall, cfDNA demonstrated significantly higher affinity toward the 8BA structure compared to the 8PA system, confirming that the enhanced hydrophobic interactions from the porous structure and hydrogen bonding from 8BA molecules contributed to the binding of cfDNA. Microscopy images of cfDNA staining revealed that 8BA crystals exhibited strong green fluorescence around the crystal edges when incubated with cfDNA, while the no-cfDNA control and 8PA with cfDNA groups showed no such phenomenon (Fig. 4k).

### 2.3. Anti-inflammatory Activity of 8BA and Mechanistic Insights

#### 2.3.1. Biocompatibility, Blood Compatibility and Pharmacokinetics of 8BA

To further explore the biomedical applicability of 8BA, its biocompatibility and blood compatibility were investigated. To observe the cytotoxic effects of 8BA materials at different concentrations on cells, cell viability was assessed through live/dead staining and CCK-8 (Cell Counting Kit-8) assays on BMM (bone marrow mononuclear) cells and RAW264.7 cells treated for 24, 48, and 72 hours. For BMM cells, a decrease in green fluorescence signals was only observed when the material concentration exceeded 1 mM after 3 days of treatment. Combined with CCK-8 results, this indicates that 8BA at approximately 1 mM exhibits negligible toxicity to BMM cells; however, with concentrations exceeding 1 mM, cell viability gradually declined as the concentration of 8BA increased (Supplementary Fig. 20). The results for RAW264.7 cells were similar to those of BMM cells, suggesting that 1 mM 8BA exhibits good biocompatibility with cells (Supplementary Fig. 21). Subsequently, the blood compatibility of 8BA at different concentrations was investigated, and the results were compared with the classical cationic adsorbent PAMAM-G3 at the same molar concentration. The results showed that, for the 8BA treatment groups at concentrations of 2 mM or lower, the supernatant was colorless, similar to the negative control PBS group, while the supernatant in the 5 mM 8BA treatment group appeared light red, indicating partial hemolysis. In contrast, as the molar concentration of PAMAM-G3 increased, the color of the supernatant deepened, with the supernatant of the 5 mM PAMAM-G3 treatment group being close to the positive control group, appearing bright red (Supplementary Fig. 22a). Quantitative hemolysis rates revealed that the hemolysis rate for 8BA at concentrations of 2 mM or lower was below 5%, further confirming its good blood compatibility (Supplementary Fig. 22b). Finally, the pharmacokinetics and in vivo metabolism of 8BA were investigated to assess its biological safety. In this experiment, 8BA at doses of 5 mg/kg and 10 mg/kg were administered to mice via intravenous injection and oral gavage, respectively. Plasma drug concentrations within 24 hours post-administration indicated the presence of the parent drug within the first hour for both administration routes, demonstrating that 8BA is more stable in vivo compared to 2FA (< 1 min). Based on the changes in plasma concentrations of 8BA over time, the main pharmacokinetic parameters were calculated, with no abnormal metabolism observed. Furthermore, compared to 2FA, 8BA exhibited a longer half-life, suggesting that 8BA has a longer *in vivo* residence time than 2FA (Supplementary Table 5-6 and Supplementary Fig. 23). Additionally, major metabolites of 8BA were identified using ultra-high performance liquid chromatography-quadrupole time-of-flight mass spectrometry (UHPLC/Q-TOF-MS/MS), and the *in vivo* metabolic profile of 8BA was elucidated. Phase I metabolic reactions of 8BA included oxidation, hydration, and desaturation, while phase II reactions involved arginine conjugation and glutathione conjugation. The primary *in vivo* metabolism of 8BA involved the cleavage of glycosidic bonds, F-deoxygenation at the 2’-position of the ribose, and reduction at the benzyl alcohol position, with no abnormal metabolites detected (Supplementary Fig. 24-25). Furthermore, histopathological examinations of the heart, liver, spleen, lungs, and kidneys revealed no organ toxicity in mice treated with 8BA, indicating the *in vivo* biological safety of 8BA (Supplementary Fig. 26).

#### 2.3.2. Transcriptomic Analysis of Anti-inflammatory Mechanism of 8BA

Previous studies demonstrated that 2FA could inhibit TLR receptor mRNA expression and osteoclast differentiation. Thus, whether 8BA retained these capabilities was further investigated. RNA sequencing results revealed that 8BA could inhibit the expression of Tlr3, Tlr4, Tlr7, Tlr9 and Tlr13. Specifically, the expression of Tlr7, Tlr13 and Tlr4 was inhibited by approximately 3-fold, while Tlr9 and Tlr3 expression was reduced by approximately 2-fold (Extended Data Fig. 3a). Gene Ontology (GO) and Gene Set Enrichment Analysis (GSEA) demonstrated that LPS stimulation significantly upregulated multiple inflammation-related pathways in cells, including cytokine production, inflammatory response, immune response, TLR signaling pathway and IL-17 signaling pathway (Extended Data Fig. 3b, d). When 1 mM 8BA was introduced simultaneously with LPS, cytokine-related pathways, immune cell differentiation pathways and osteoclast differentiation pathways were downregulated, while bone generation pathways were upregulated (Extended Data Fig. 3c, e). A schematic illustration showed that 8BA could adsorb LPS and cfDNA while simultaneously inhibiting the expression of Tlr3, Tlr4 and Tlr9, thereby suppressing the transcription of downstream inflammatory factors (Extended Data Fig. 3f).

Quantitative real-time PCR (qRT-PCR) validation confirmed that under LPS stimulation, 1 mM 8BA downregulated Tlr4 expression by more than 2-fold. Under cfDNA stimulation, 1 mM 8BA reduced Tlr3 expression to approximately 60% and Tlr9 expression to approximately 50% of the stimulated levels (Extended Data Fig. 3g). Additionally, 8BA retained the ability of 2FA to inhibit osteoclast differentiation, while showing no significant effect on osteoblast differentiation. After 7 days of co-culture of different concentrations of 8BA with BMM cells under osteoclast induction, TRAP (tartrate-resistant acid phosphatase) staining results indicated that 1 mM 8BA could inhibit osteoclast differentiation (Extended Data Fig. 3h). After 7 days of co-culture of different concentrations of 8BA with BMSC (bone marrow mesenchymal stem cells) cells under osteoblast induction, ALP (alkaline phosphatase) staining results showed that 8BA had no significant effect on osteoblast differentiation (Supplementary Fig. 27). Western blot (WB) was performed to detect the expression levels of key proteins in TLR4, TLR9, MAPK and NF-κB signaling pathways. The results demonstrated that 8BA could effectively inhibit the excessive inflammatory response induced by LPS and cfDNA at the protein level (Extended Data Fig. 3i). Immunofluorescence staining results indicated that the protein expression levels of TLR4 and TLR9 were reduced to less than 50% of the inflammatory levels (Extended Data Fig. 3j).

#### 2.3.3. *In vivo* Anti-inflammatory Effects of 8BA in Periodontitis and Psoriasis Models

To investigate whether 8BA could alleviate tissue inflammation by binding pro-inflammatory factors such as cfDNA and LPS, periodontitis models were firstly established. Mouse periodontal tissues were collected and subjected to inflammation level analysis and Micro-CT examination. In the early treatment of periodontitis (Fig. 5a), the 8BA treatment groups exhibited lower salivary TNF-α, IL-6 and cfDNA levels compared to other material treatment groups, with levels significantly lower than other groups and closer to those of healthy periodontal tissue. Immunohistochemical (IHC) staining revealed that the 8BA treatment groups showed lower positive rates of TLR4, TLR9, IL-6 and TNF-α in periodontal tissues, reaching approximately 40% of the periodontitis levels, suggesting that 8BA could effectively alleviate periodontal tissue inflammation (Fig. 5b-c and Supplementary Fig. 28). In the late treatment of periodontitis (Extended Data Fig. 4), the 8BA groups showed lower distance from the cement-enamel junction to the alveolar crest, and higher buccal-lingual bone density. H&E and MESSON staining results demonstrated that the 8BA treatment groups exhibited lower periodontal pocket depth compared to other treatment groups. TRAP staining results indicated fewer osteoclasts in the 8BA groups compared to other groups, further suggesting that 8BA could promote hard tissue repair by inhibiting osteoclast formation (Extended Data Fig. 4).

**Fig. 5.**
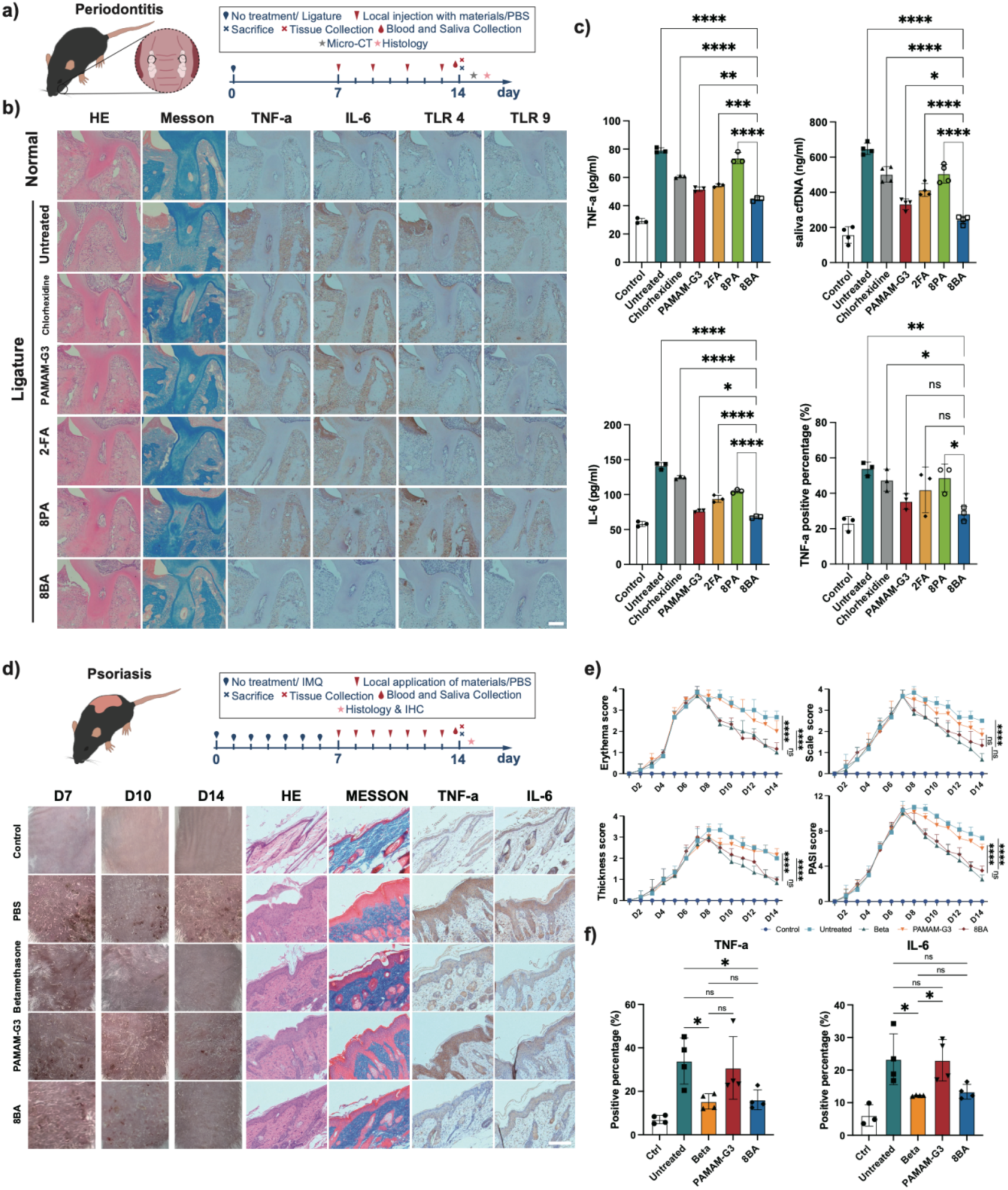
Local administration of 8BA effectively alleviates periodontitis and psoriasis. a) Schematic illustration showing the establishment of periodontitis model. b) Representative images of H&E, MESSON, TNF-α, IL-6, TLR4 and TLR9 IHC staining in tooth and periodontal tissues of non-periodontitis, periodontitis, and periodontitis groups treated with chlorhexidine, PAMAM-G3, 2FA, 8PA or 8BA. c) Bar plot showing TNF-α, cfDNA and IL-6 levels in saliva of mice in different groups, and the positive rate of TNF-α in IHC staining. n = 3. d) Schematic illustration showing the establishment of psoriasis model. Lower panel shows control and psoriasis groups treated with PBS, Betamethasone, PAMAM-G3 or 8BA. Representative images of H&E, MESSON, TNF-α and IL-6 IHC staining. e) Line plot showing Erythema score, Scale score, Thickness score and PASI score in different groups over time, n = 4. f) Bar plot showing TNF-α and IL-6 IHC positive ratio in different groups, n = 4. Statistical analyses were performed using one-way ANOVA with Tukey’s multiple comparisons test in (c and f). In (c and f), bar graphs are shown as mean ± SD. * p < 0.05, ** p < 0.01, *** p < 0.001, **** p < 0.0001. Scale Bar = 200 µm.

The therapeutic effect of 8BA was further evaluated in a mouse model induced by imiquimod (IMQ), which served as an acute psoriasis-like mouse model as it could induce skin inflammatory responses resembling certain aspects of human psoriasis. As a TLR7/8 agonist, IMQ could induce a series of skin reactions, leading to erythema, scaling and skin plaque thickening, indicating psoriasis onset (Fig. 5d). Throughout the experimental period, IMQ-induced mice showed significantly higher Psoriasis Area and Severity Index (PASI) scores compared to normal mice. During the following 7 days, betamethasone, PAMAM-G3 and 8BA were applied topically to the psoriatic skin of IMQ mice. During the treatment period, 8BA demonstrated superior therapeutic effects compared to PAMAM-G3, as evidenced by reduced scaling and erythema, and PASI scores significantly lower than the model group and approaching those of the betamethasone group (Fig. 5e). In the TNF-α and IL-6 IHC positive ratio statistics, the levels of both markers in the 8BA group were essentially similar to those in the betamethasone group, while the expression levels in the PAMAM-G3 group were approximately 2-fold higher than those in the 8BA treatment group (Fig. 5f).

## 3. Discussion

In this study, an SE(3)-equivariant deep-learning workflow, SE3CSP, is developed to enable prospective crystal structure prediction and structure-guided design of nucleoside self-assembling materials. Notably, in recent years, CSP has evolved from a thermodynamic search for global lattice-energy minima into a broader framework for mapping the structural landscape of molecular solids, yet remains fundamentally constrained by the coupled challenges of exhaustive sampling and reliable candidate prioritization, particularly for flexible molecules^17,22,67^. Classical studies have shown that crystal structures are governed by subtle balances of intermolecular interactions, and that experimentally realized forms do not necessarily correspond to the global minimum, as highlighted by successive CSP blind tests^12^. These limitations arise in part because conformational flexibility couples intramolecular and packing degrees of freedom, making it difficult to generate relevant conformers and packing motifs during early-stage sampling, such that correct structures may be excluded before ranking^18,68,69^. Recent advances have addressed this challenge along two directions: improving ranking accuracy through dispersion-corrected DFT and machine-learning interatomic potentials, and reshaping structural generation via data-driven approaches^21,70,71^. In particular, emerging deep-learning frameworks, including generative and diffusion-based models trained on experimental crystal structures, have demonstrated improved sampling of realistic conformations and packing arrangements (e.g., DeepCSP, OXtal), yet still rely on physically grounded refinement for validation^23,33^. Within this context, SE3CSP integrates SE(3)-equivariant learning of molecular conformations, unit-cell parameters and packing patterns with MACE-based optimization, effectively coupling learned structural priors with efficient energy refinement. Unlike workflows that primarily accelerate post-generation ranking, SE3CSP shifts the predictive burden upstream by improving the quality of candidate structures entering refinement, which is critical for flexible nucleosides and related self-assembling systems. The incorporation of a density-based pre-filter, followed by energy ranking and structural deduplication, further converts a broad CSP landscape into a tractable prospective search space, enabling the experimentally closest structures to be consistently prioritized within the top 1-2% of generated candidates (Table 1). Together, these results indicate that SE3CSP not only improves predictive accuracy, but redefines CSP as a more prospectively actionable framework for structure-guided materials design.

Chemical modification of organic self-assembling molecules has long been a powerful strategy for generating new supramolecular functions, because subtle changes in hydrogen-bonding motifs, aromatic surfaces, hydrophilic–hydrophobic balance and molecular conformation can markedly reshape assembly pathways and material properties^72,73^. Nucleosides are particularly attractive building blocks in this context owing to their intrinsic biocompatibility, structural programmability, multiple hydrogen-bond donor and acceptor sites, and capacity to assemble through coupled hydrogen bonding, π–π stacking and hydrophobic interactions. These features have supported the development of diverse nucleoside- and nucleobase-derived supramolecular systems, including hydrogels, hierarchical assemblies and biofunctional materials^74-76^. In our previous studies, 2FA was found to possess strong self-assembly capability, and targeted molecular modification provided an effective means to regulate its crystal packing and supramolecular morphology, highlighting crystallographic insight as a foundation for nucleoside material design^45,46^. In the present work, SE3CSP extends this principle by enabling prospective evaluation of how molecular modification alters conformational preference, packing motifs and accessible energy landscapes, thereby providing a computational route to guide nucleoside self-assembly before exhaustive experimental screening. More importantly, the CSP-guided design of 8BA yielded an adsorptive nucleoside material that departs from conventional inflammation-scavenging adsorbents, many of which rely predominantly on cationic charge to capture negatively charged inflammatory mediators such as cfDNA and LPS^77^. Although electrostatic adsorption is effective, strongly cationic materials can also induce nonspecific biomolecular interactions, membrane perturbation and toxicity, which remain important concerns for biomedical use^78-80^. By contrast, 8BA integrates abundant hydrogen-bond donor/acceptor groups with a dynamic breathing porous structure, allowing inflammatory mediator adsorption through cooperative non-electrostatic interactions, including hydrogen bonding, π-π stacking and confinement-assisted uptake (Fig. 4f). This design is consistent with the broader concept of soft porous crystals, in which adaptive pore motion can facilitate guest adsorption and structural responsiveness. Thus, SE3CSP-guided nucleoside modification not only expands the structural design space of bioactive self-assembled materials, but also offers a route to overcome the charge–biocompatibility trade-off that has limited many positively charged adsorbents.

In overall, these results establish SE3CSP as a generalizable workflow for organic crystal structure prediction and demonstrate its utility in nucleoside self-assembled materials as a representative testbed. The improved performance on CSD blind-test candidate molecules suggests that the model is not restricted to nucleosides, and future studies may extend this framework to broader classes of flexible organic solids. This expansion is particularly timely because recent CSP blind tests continue to show that molecular complexity, conformational flexibility, multicomponent crystallization and polymorphism remain major frontiers for the field. At the same time, most current CSP and AI-assisted CSP workflows, including the present implementation, are still primarily optimized for single-conformer, solvent-free crystals. Extending SE3CSP to crystals containing multiple independent conformers, disordered components, solvates, hydrates or host–guest architectures would be especially valuable, because solvent and guest positions often reshape the packing landscape and can stabilize conformations that differ from those found in unsolvated forms. Deep learning may offer a useful route to address this problem because equivariant architectures can, in principle, learn correlated molecular conformations, local environments and periodic host–guest geometries directly from experimental structural data, rather than treating solvent placement as a purely post hoc packing problem. The rapid growth of molecular-crystal datasets and foundation-model efforts, such as OMC25, may further support this direction by providing larger and more diverse training spaces for crystal environments beyond simple Z′ = 1 structures^81^. Another important direction is the modular integration of SE3CSP with different physical refinement engines. In this work, MACE-off was used as the downstream optimizer because machine-learning force fields have recently shown strong promise for organic molecules and molecular crystals, providing a balance between transferable accuracy and computational efficiency. However, different force fields and machine-learning interatomic potentials may differ in their treatment of dispersion, polarization, hydrogen bonding, conformational strain and finite-temperature effects, all of which are central to molecular crystals. Recent studies on MLIPs for molecular crystals and CSP indicate that active learning, batched optimization and system-specific potentials may further improve the reliability of ranking and relaxation^71,82^. Thus, a key future question is not only which force field is most accurate in isolation, but which refinement model is best matched to the structures generated by the preceding deep-learning stages. Addressing this coupling between learned generation and physical refinement will be essential for transforming SE3CSP from a successful workflow for nucleoside assemblies into a broadly deployable platform for prospective molecular crystal design.

## 4. Methods

### 4.1. The Development of SE3CSP

We present SE3CSP, an SE(3)-equivariant deep learning model for crystal structure prediction (CSP) of organic molecular crystals. Given a molecular simplified molecular input line entry system (SMILES) string as the sole input, SE3CSP simultaneously predicts the crystalline conformation of the molecule and the corresponding lattice parameters. The model is built upon the Geometric Algebra Transformer (GATr) framework, which naturally encodes three-dimensional (3D) spatial information through multivector representations in the geometric algebra of 3D Euclidean space, thereby achieving full rotational and translational equivariance by construction. The overall architecture comprises three cascaded modules, including a conformational packing module (Fig. 1a), a lattice prediction module (Fig. 1b), and a crystal optimization module (Fig. 1c), organized within a two-stage training pipeline (Fig. 1). Following crystal optimization, a density-guided prospective ranking strategy was further incorporated, in which structures within a predicted density window (ρ_predict_ ± 0.15 g/cm³) were retained for subsequent lattice-energy ranking and structural deduplication, enabling efficient candidate prioritization within the generated crystal energy landscape (Fig. 1d).

### 4.2. Problem Definition

Crystal structure prediction requires determining both molecular-level geometry and crystal-level periodicity from minimal input. Crystal structures encode two coupled sources of information: the conformational geometry of individual molecules, and the periodic lattice that packs these molecules into a repeating solid. Searching through all possible packing configurations for even a moderately sized organic molecule requires exploring an astronomically large conformational space, in which minor perturbations of atomic positions can produce qualitatively different packing arrangements and vastly different material properties.

SE3CSP reframes organic molecule CSP as a learned prediction task. Given an input molecule encoded as a SMILES string, the model must determine both the 3D molecular conformation within the crystal lattice and the crystallographic lattice parameters that govern the periodic packing. Formally, let *S* denote the SMILES string and 𝒞^In^ the initial 3D conformation generated from *S*, SE3CSP models the conditional probability distribution:

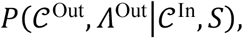

where 𝒞^Out^ denotes the predicted 3D molecular conformation packed inside the crystal lattice, and 𝛬^Out^ represents the crystallographic parameters including lattice lengths (*a*, *b*, *c*), lattice angles (𝛼, 𝛽, 𝛾), the space group, and the crystal density 𝜌.

This problem is decomposed into two coupled subtasks. The first is predicting the 3D molecular conformation, which establishes the relative positions of all atoms. The second is determining the crystallographic unit cell parameters, which dictate how the molecule replicates to fill 3D space. These subtasks are inherently linked: the molecular conformation determines which packing arrangements are energetically favorable, while the symmetry operations imposed by the space group restrict the allowable conformations. The overall architecture is illustrated in Fig. 1.

### 4.3. Dual Sequence Molecular Representation

Standard SMILES notation encodes only the 2D molecular graph and treats the molecule as a single unfragmented string, discarding local chemical environment information that strongly influences conformational preferences in the crystalline state. To overcome these limitations, SE3CSP adopted a dual sequence-based SMILES representation syntax proposed in our previous study, named DSMILES.

In DSMILES, the input molecule *S* is first decomposed into chemically meaningful fragments (such as individual ring systems and their substituents), and each fragment is encoded as a sub-SMILES string. Two sequences of equal length are then produced. The first is the DSMILES token sequence {𝑡_1_, 𝑡_2_, …, 𝑡_𝐿_} of maximum length 𝐿, where each token carries annotations for element type and ring-system classification. The second is a positional pointer sequence {𝑝_1_, 𝑝_2_, …, 𝑝_𝐿_} of the same length, where each entry specifies the index of the atom to which the current token connects, thereby encoding inter-fragment connectivity. An illustrative example of the DSMILES encoding for a sample molecule alongside its canonical SMILES is provided in Extended Data Figure 1.

Once the dual sequences are generated from the SMILES string, the corresponding conformational coordinates are also arranged into a sequence of the same length. Each entry contains the atomic coordinates, with non-element positions set to zero.

### 4.4. SE(3)-Equivariant Geometric Attention Backbone

A central design requirement for any deep learning model processing molecular and crystallographic data is that it must respect the fundamental symmetries of 3D space. Specifically, the predicted molecular conformation must transform identically to the input under any rotation or translation, necessitating that the model be SE(3)-equivariant. SE3CSP adopts the GATr architecture^83^, which enforces SE(3)-equivariance by construction through multivector representations grounded in Clifford algebra. This guarantees that every operation within the network, from attention computation to nonlinear activations, preserves geometric structure.

The GATr encoder consists of multiple equivariant transformer blocks. Each block sequentially applies equivariant layer normalization, geometric self-attention with rotary encodings, and a geometric MLP (GeoMLP) that uses geometric products and joins to nonlinearly transform multivector features. Residual connections are included after each major operation. Rotary encodings extend the rotary position encoding (RoPE) mechanism to multivectors by applying grade-specific rotations to scalar, vector, bivector, and trivector components. This ensures the encoding remains equivariant when both keys and queries are rotated together, making it particularly effective for processing geometric and spatial data in transformer models. The encoder outputs updated multivectors, scalars, and a global reference multivector that defines the equivariant frame. Architectural details are provided in Supplementary Material Section 1.1.

### 4.5. Conformational Packing Module

The conformational packing module is the first of three cascaded modules in SE3CSP (Fig. 1a). It predicts the crystal packing conformation of the input molecule, which then serves as the geometric input for the lattice prediction module. This module comprises an embedding layer, a GATr encoder, and three specialized prediction heads.

#### 4.5.1. Embedding Layer

Given an input molecule encoded as a SMILES string *S*, an initial 3D conformation 𝒞^In^ is generated using the RDKit package, producing a reasonable force-field level starting geometry. Subsequently, both *S* and 𝒞^In^ are represented using the DSMILES syntax. The input to the embedding layer consists of three components: the DSMILES token sequence, the positional pointer sequence, and the initial 3D atomic coordinates.

The GATr architecture processes input through dual parallel streams: one for scalars and another for multivectors. To represent the scalar embedding of each token, the model first projects the token through four parallel embedding pathways. These embeddings are obtained from the DSMILES token sequence and the positional pointer sequence, and are subsequently combined via element-wise summation:

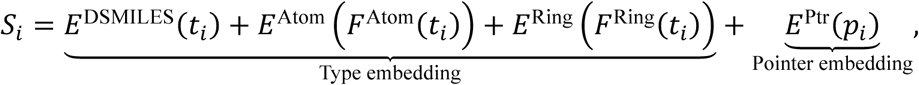

where 𝐸^DSMILES^ is the DSMILES token embedding, *E*^Atom^ maps the element type extracted from the DSMILES token via 𝐹^Atom^, *E*^Ring^ maps the ring system classification extracted from the DSMILES token via 𝐹^Ring^, and 𝐸^Ptr^ is the positional pointer embedding. Using decomposed atom and ring embeddings, rather than a single DSMILES token embedding, provides an inductive bias that encourages sharing of representations across tokens with the same element type or ring system, improving generalization to rare token types.

Furthermore, the multivector embedding of each token is constructed by representing its 3D coordinate triplet 𝑟_𝑖_ = (𝑥_𝑖_, 𝑦_𝑖_, 𝑧_𝑖_) as a geometric algebra multivector. This is achieved via a trivector formulation that maps Cartesian coordinates into the conformal geometric algebra Cl(3,0,1):

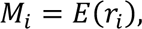

where each coordinate triplet is projected into a single multivector channel utilizing the trivector embedding formula described in the Supplementary Material Section 1.1.1.

The above explicit separation of scalar embeddings encoding chemical and topological information from multivector embeddings encoding geometric information is fundamental to the dual-stream architecture of the GATr backbone. It enables the two streams to evolve independently while interacting through a geometric self-attention mechanism.

#### 4.5.2. Conformational Encoder

The embedded multivectors and scalars from the embedding layer are processed by the conformational encoder. This encoder follows the GATr architecture (Section 2.3), which comprises multiple equivariant transformer blocks. After processing through the encoder, the model produces a refined latent representation for the scalar stream *S*^𝒞^, a latent representation for the multivector stream 𝑀^𝒞^, and a reference multivector 𝑅^𝒞^ for the equivariant join operation. Further implementation details are provided in Supplementary Material Section 1.1.2.

#### 4.5.3. Auxiliary Prediction Heads

The conformational encoder generates rich geometric and chemical representations for each atom, which are subsequently decoded into concrete predictions through three specialized GeoMLP-based prediction heads. Among these, the token prediction head and the pointer prediction head serve as auxiliary components during training, designed to enhance the model’s representational consistency and topological awareness.

##### Token Prediction Head

This head predicts the DSMILES token type for each position, outputting a probability distribution over the DSMILES vocabulary:

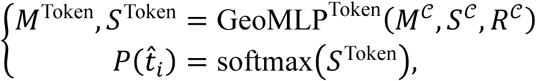

It acts as an auxiliary supervision signal, ensuring that the model preserves molecular identity throughout the conformation prediction process.

##### Pointer Prediction Head

Similarly, the pointer prediction head is not used during inference because the molecular topology is already known at that phase. However, it aids learning in the training phase by predicting positional pointers between atoms conditioned on the corresponding token types:

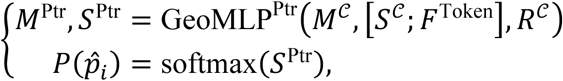

where [*S*;*F*^token^] denotes the concatenation of the GATr scalar output and the token embedding along the feature dimension. This head reinforces topological consistency during conformation prediction.

#### 4.5.4. Position Prediction Head

Following a design similar to that of the pointer prediction head, the position prediction head estimates the 3D atomic coordinates within the crystalline environment, conditioned on both DSMILES token types and positional pointers:

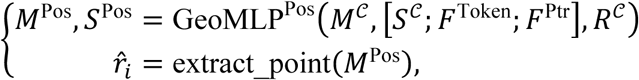

where 𝑀^Pos^ is projected to three Cartesian coordinates, and ‘extract_point’ denotes the operation that extracts the 3D atomic coordinates for each atom from its corresponding 16D multivector representation.

Although the predicted coordinates 𝑟_𝑖_ are global, local geometric properties, including bond lengths, bond angles, and dihedral angles, are computed from them. These properties serve as auxiliary geometric supervision signals that encourage the model to learn chemically meaningful local geometries. More details about loss functions are provided in Supplementary Materials Section 1.2.1.

### 4.6. Lattice Prediction Module

While the conformational packing module determines internal molecular geometry, crystal structure prediction additionally requires determining how molecules are arranged in 3D periodic space. The lattice prediction module extends SE3CSP with a dedicated secondary GATr backbone and three specialized prediction heads, designed to learn the mapping from molecular spatial arrangement to crystallographic parameters.

#### 4.6.1. Lattice Encoder

The lattice encoder is a shallower GATr encoder that processes the predicted conformer coordinates, which are embedded as multivectors, alongside the original scalar embeddings. This architectural separation serves two purposes. First, crystallographic parameters depend primarily on the global spatial envelope of the molecule, such as its overall shape, size, and symmetry, rather than on fine-grained atomic details; therefore, a shallower network is sufficient. Second, the separation enables the selective parameter-freezing strategy, where the primary backbone remains frozen while the lattice backbone is trained. Finally, the lattice encoder produces lattice multivectors *M*^ℒ^, scalars *S*^ℒ^, and reference multivector *R*^ℒ^.

#### 4.6.2. Lattice Prediction Heads

Building on the output of the lattice encoder, we design two specialized prediction heads based on GeoMLP to learn the molecular envelope.

##### Space Group Prediction Head

The space group prediction head processes a learned first token derived from the lattice encoder output. This special token, which is prepended to the input sequence, begins with a randomly initialized embedding that is updated via self-attention to aggregate global sequence information. Subsequently, the head transforms this representation through a GeoMLP to produce logits over all supported space groups:

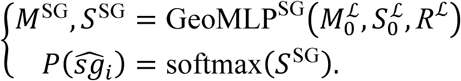

These supported space groups were selected based on frequency analysis of the CSD, and account for approximately 90% of organic molecular crystal entries meeting these criteria. The predicted space group then conditions the lattice parameter prediction through a learnable embedding, as the symmetry constraints of different space groups impose distinct relationships among the lattice parameters.

##### Lattice Parameter Prediction Head

Given the predicted space group 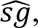 the lattice parameter head predicts the six lattice parameters (*a*, *b*, *c*, 𝛼, 𝛽, 𝛾) along with the crystal density 𝜌. This head processes the representation of the first token from the lattice encoder, conditioned on the predicted space group via a learnable space group embedding that is concatenated with the scalar features:

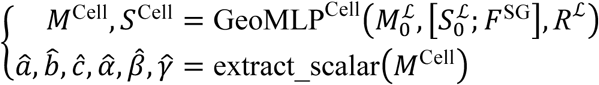

Here, 𝐹^SG^ is the embedding of the predicted (or ground-truth during training) space group. The lattice parameters are extracted as scalar components from the output multivectors. Additionally, the crystal density *ρ* is predicted by a parallel GeoMLP density projection head. In addition, the details regarding the loss functions used in the lattice prediction module are provided in Supplementary Materials Section 1.2.2.

### 4.7. Crystal Optimization Module

The conformational packing and lattice prediction modules generate preliminary crystal structures whose atomic coordinates and lattice parameters are obtained from neural network predictions rather than physical energy minimization. To refine these structures into physically realistic local minima on the potential energy surface, we apply a force-field optimization module (Fig. 1c) based on MACE-OFF. This module acts as a deterministic post-processing step with no feedback loop to the neural network, thereby ensuring that the preceding modules remain unbiased by any specific force field. For each target molecule, approximately 2,500 preliminary structures are subjected to force-field optimization; the resulting energies and geometries are then used to construct the crystal energy landscape for subsequent analysis.

Following the approach described in a previous study^84^, we implement a two-stage optimization protocol that mirrors common practice in crystal structure prediction. In the first stage, an external hydrostatic pressure is applied while both atomic positions and lattice parameters are optimized simultaneously. This pressure term drives the unit cell toward a physically realistic density, steering the structure away from artificially expanded configurations that can arise from neural network predictions. In the second stage, the external pressure is removed, and the structure undergoes a final energy minimization at zero pressure to converge to the true local minimum.

### 4.8. Structure Evaluation, Redundancy Reduction, and Ranking Method

To rigorously validate the predictive capability of SE3CSP, the crystal structures generated by the above three modules were systematically benchmarked against the experimental reference using the COMPACK algorithm (implemented within the CCDC Mercury suite)^85^. A prediction is considered successful if a minimum of 8 molecules within a 15-molecule packing shell align with the reference structure, and the root-mean-square deviation (RMSD₁₅) is below 2.0 Å.

#### 4.8.1. Redundancy Reduction

Due to the continuous nature of the potential energy surface, generated structures often contain near-duplicate structures that can obscure the true distribution of packing landscapes. To address this, we implement a rigorous structural deduplication method based on molecular overlay similarity. By performing pairwise comparisons within the 15-molecule coordination shell, we identify structures that share 12 or more matched molecules, indicating a similarity threshold of 80% or higher. These highly similar structures are treated as redundant entries representing the same packing motif; duplicates are pruned to ensure that only unique, representative crystal packings are retained for final analysis.

#### 4.8.2. Final Crystal Ranking Methods

After filtering for structural uniqueness, the final step involves identifying the most thermodynamically plausible structure by navigating the crystal energy landscape. We employed two distinct ranking strategies:

##### Solely Energy-Based Ranking

Predicted crystal structures were ranked exclusively according to their lattice energies after MACE-OFF optimization, following the conventional thermodynamic assumption in CSP that the experimentally realized structure corresponds to, or lies close to, the global minimum on the crystal energy landscape^8,44^. However, purely energy-based ranking remains challenging for flexible molecular crystals because the energy landscape is highly multidimensional and contains numerous competing local minima. In addition, lattice energy alone does not account for uncertainties arising from force-field inaccuracies, conformational variability or non-physical prediction artifacts, which can occasionally lead to misleadingly favorable low-energy structures^30,86,87^. Consequently, non-physical artifacts, such as the artificially expanded unit cells that arise from neural network predictions (Section 4.7), may yield misleadingly low energies and obscure the true global minimum.

##### Density-Guided Prospective Ranking

To reduce the limitations associated with purely energy-based ranking, a density-guided filtering strategy was introduced prior to lattice-energy ranking. Specifically, only predicted structures with densities falling within a tolerance window of ρ_predict_ ± ρ_T_ g/cm³ were retained for subsequent energy evaluation. By incorporating predicted crystal density as an additional physical constraint, this approach effectively removes structurally unrealistic candidates before ranking, thereby improving the likelihood of identifying experimentally relevant low-energy structures. A series of benchmark analyses was further performed to optimize the density tolerance parameter (ρ_T_), and the results showed that ρ_T_ = 0.15 provided the best overall predictive performance among the tested thresholds (Supplementary Table 3).

### 4.9. Two-stage Data-driven Training Strategy

Crystal structure prediction requires a model to simultaneously capture atomic-level molecular geometry and crystal-level periodicity, a dual objective that introduces distinct learning challenges at different scales of structural organization. Training a single model end-to-end to predict both molecular conformations and unit cell parameters risks instability, as the two tasks operate on different feature spaces and demand representations of differing complexity. To address this, SE3CSP adopts a sequential two-stage training strategy that progressively advances from learning molecular conformations to predicting crystal-level structures. This approach first learns to accurately reproduce molecular geometry, then extends the acquired geometric representation to crystallographic prediction. This design reflects the natural hierarchical organization of crystal structures and establishes a stable foundation for the more complex task of lattice parameter learning in the second stage.

#### 4.9.1. Data Collection and Inclusion Criteria

The data used in this study were obtained from the Cambridge Structural Database (CSD), which encompasses a wide variety of crystal types, including both organic and inorganic structures, as well as solvated and solvent-free forms. This diversity presents considerable challenges for model training. To mitigate the complexity of the learning task, we applied the following inclusion criteria: (a) only organic crystals were retained; (b) structures were required to be single-conformation (Z’=1) and solvent-free; and (c) the SMILES representation of each crystal had to be encodable using the DSMILES syntax^88^. According to the above inclusion criteria, a total of 12 molecules from previous CSD blind-test candidates were retained for evaluation of the model performance.

#### 4.9.2. Conformation Learning Stage

The first training stage is dedicated solely to the molecular conformation prediction task, establishing the foundational geometric representations upon which the model depends in all subsequent stages. The core learning objective is to train the GATr backbone and all three prediction heads (token, pointer, and position) to accurately reconstruct experimentally determined crystal conformations from noisy initial geometries.

Each training sample comprises a pair of molecular conformations: an input conformation generated by the RDKit package via a force-field-based conformer generator, which produces chemically reasonable but crystallographically approximate geometries, and a target conformation taken directly from the experimental crystal structure. This input-target pairing embodies the fundamental learning signal: the model must learn to refine approximate molecular geometries into crystal-accurate conformations.

#### 4.9.3. Crystal Parameter Learning Stage

Having established robust molecular conformation representations in the first learning stage, the second training stage extends SE3CSP to predict crystallographic lattice parameters. This stage incorporates a dedicated lattice prediction module, which consists of a separate lattice encoder together with specialized prediction heads for space group classification, lattice parameter regression, and crystal density estimation. The module leverages the pre-trained molecular geometry knowledge obtained from the conformation learning stage.

To preserve the previously acquired molecular geometry while training the newly introduced lattice module, all parameters of the pre-trained conformational encoder and its three prediction heads are kept frozen. Only the lattice encoder and its corresponding prediction heads are updated during the crystal parameter learning stage. This selective parameter freezing strategy is critical: updating the conformation backbone could cause catastrophic forgetting of the geometric representations that form the foundation of the entire system, whereas training only the unit cell module allows it to learn a mapping from molecular geometry to crystallographic parameters without corrupting the underlying geometric understanding.

### 4.10. Materials, Cells and Animals

All chemicals were commercially sourced and used without further purification unless otherwise specified. 2-Amino-2′-fluoro-2′-deoxyadenosine (2FA, CAS: 134444-47-6) was obtained from Shanghai Yuanye Biotechnology Co., Ltd (purity > 99%) and Wuhu Nuowei Chemistry Co., Ltd (purity > 97%). Other reagents were purchased from Sigma-Aldrich (Shanghai, China), J&K, Adamas, Acros Organics, Bide Pharmatech, Chengdu RZBT Chemical Reagent Co., Ltd, or Energy Chemical. Ultrapure water used in all experiments was generated using a Milli-Q Plus system. The synthetic routes of the 8-position derivatives of 2FA are shown in Supplementary Fig. 1.

RAW 264.7 cells were cultured in Dulbecco’s modified Eagle’s medium (DMEM, Gibco) supplemented with 10% fetal bovine serum (FBS) at 37 °C in a humidified atmosphere containing 5% CO₂. For BMSC and BMM isolation, mouse femurs were harvested under sterile conditions, and bone marrow was flushed out with phosphate-buffered saline (PBS) using a 27-gauge needle. After filtration through a 70 μm strainer (Corning), BMMs were isolated using CD11b-positive magnetic beads (Miltenyi Biotec) according to the manufacturer’s instructions. BMSCs were cultured in α-MEM (Gibco) containing 10% FBS, 1% penicillin–streptomycin and 1% L-glutamine. Non-adherent cells were removed after 24–48 h, and cells at passages 3–5 were used for subsequent experiments. Cell viability was assessed by trypan blue staining before use or cryopreservation.

Male C57BL/6 mice (8–9 weeks old) and female BALB/c mice (8–9 weeks old) were purchased from Chengdu Dake Experimental Animal Co., Ltd and housed under specific pathogen-free conditions at the Experimental Animal Center of Sichuan University with free access to food and water. Animals were acclimatized for at least one week before experiments.

### 4.11. *In Vitro* Investigation of the Adsorption Behavior and Underlying Mechanism

#### 4.11.1. Binding Ability Assessment

To test cfDNA binding ability, EtBr competition array was performed. 10 μL of DNA (20 μg/mL) and 10 μL of EtBr (20 μg/mL) were added to 170 μL of PBS in a 96-well plate. Then, 10 μL of PAMAM-G3, 2FA, chlorhexidine (at a fixed concentration), 8PA, and 8BA at varying concentrations were added to each well. Fluorescence intensity was measured using a plate reader at 595 nm. The nucleic acid binding affinity (X) was calculated using the formula:

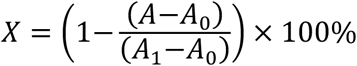

Where X represents the nucleic acid binding affinity, A is the fluorescence intensity of the sample, A0 is the fluorescence intensity of EtBr without DNA and polymers, and A1 is the fluorescence intensity of the EtBr/DNA complex without polymers.

For binding ability of LPS, TNF-α and IL-6, PAMAM-G3, 2FA, chlorhexidine (at a fixed concentration), 8PA, and 8BA at varying concentrations incubated in LPS solution (1 mL, 10 ng/mL), TNF-α solution (1 mL, 10 ng/mL), and IL-6 solution (1mL, 200 pg/mL) respectively, while shaking (60 rpm) at 25 °C for 12 h. Then, the supernatants were obtained and ELISA kits were used to measure the TNF-α/LPS/IL-6 levels.

#### 4.11.2. TNF-α and IL-6 Adsorption Performance in Cell Supernatant

Cells were seeded in 6-well plates at a density of 2 × 10⁵ cells per well and treated with 100 ng/mL LPS or 100 ng/mL cfDNA, as well as 1 mM and 2 mM 8BA for 24 hours. After treatment, cell supernatant was collected and centrifuged at 1,000 × g for 5 min to remove cell debris. The concentrations of tumor necrosis factor-alpha (TNF-α) and interleukin-6 (IL-6) in the supernatant were measured using commercially available ELISA kits (R&D Systems, Minneapolis, MN, USA) according to the manufacturer’s instructions. Absorbance was measured at 450 nm using a microplate reader (Synergy H1, BioTek Instruments, Winooski, VT, USA), and cytokine concentrations were calculated based on standard curves.

#### 4.11.3. Visualization of cfDNA Binding Efficiency Using Fluorescence Microscopy

At room temperature, cfDNA was mixed with the detection solution from the Beyotime Cell Apoptosis Kit for labeling extracellular DNA. After 2 hours, 10 µL of green fluorescent-labeled cfDNA (100 ng/mL) was added to 1 mM 8BA or 1 mM 8PA, while shaking (60 rpm) at 25 °C for 2 hours. Subsequently, the samples were fixed onto cell coverslips and imaged under a fluorescence microscope.

### 4.12. Animal Experiment

#### 4.12.1. Ligature-Induced Mouse Periodontitis Model

After one week of adaptive feeding, the mice were anesthetized via intraperitoneal injection of sterile avertin (tribromoethanol: 200 mg/10 mL kg⁻¹, dissolved in deionized water). To establish an experimental periodontitis model, a 5-0 ligature was placed at the gingival sulcus of the upper second molar, and the ligature was checked daily to ensure it remained in place.

Mice were randomly divided into seven groups: (1) healthy control group, (2) ligature + PBS treatment group, (3) ligature + chlorhexidine treatment group, (4) ligature + PAMAM-G3 treatment group, (5) ligature + 2FA treatment group, (6) ligature + 8PA treatment group, and (7) ligature + 8BA treatment group. For the experimental periodontitis model, the corresponding material (5 μL per tooth) was applied to the ligature site of the upper second molars in non-PBS treatment groups. To ensure accurate drug administration and avoid potential mucosal damage, gas anesthesia (isoflurane) was used during treatment. The ligatured mice were treated every two days. For the early periodontitis group, treatment continued until day 14, while for the late periodontitis group, treatment lasted until day 21. After saliva collection, the mice were euthanized by CO₂ inhalation, and the upper jaw was harvested for micro-CT scanning, histological staining, and molecular biology analysis.

#### 4.12.2. IMQ-Induced Mouse Psoriasis Model

BALB/c mice were anesthetized with 1.5-2% isoflurane during the experiment. To establish a psoriasis-like mouse model, a 2 cm × 2 cm area of the back skin of the mice was shaved. Then, the shaved area was treated daily with a topical dose of 62.5 mg Aldara ointment (containing 3.125 mg imiquimod, 5% IMQ; 3M Pharmaceuticals, St. Paul, MN, USA) for 7 consecutive days. After 7 days, psoriasis-like skin inflammation was successfully induced, characterized by erythema, scaling, and thickening, indicating the onset of psoriasis.

The IMQ-treated mice were randomly divided into a model group and treatment groups, with 3 mice per group. An additional 3 non-IMQ-treated shaved mice were used as the normal control group. The treatment groups included: (1) betamethasone group (positive control), (2) PAMAM-G3 group, (3) 8PA group, and (4) 8BA group. From day 8 to day 14, approximately 50 mg of 0.05% betamethasone ointment (Sigma-Aldrich) and 500 μL of the test material were topically applied to the back skin of the mice in the respective groups, followed by fixation with gauze and bandages. For the model group, 500 μL of PBS was applied to the back skin of each IMQ mouse.

The severity of skin inflammation was scored daily using an objective scoring system based on the clinical PASI (Psoriasis Area and Severity Index) scale. Erythema, scaling, and thickening were independently scored on a scale from 0 to 4: 0, none; 1, mild; 2, moderate; 3, marked; 4, very marked. The cumulative score for erythema, scaling, and thickening (0 to 12 points) was used as a measure of inflammation severity. In addition, the back skin of mice from each group was photographed on day 8 (when psoriasis symptoms were most prominent), day 11, and day 14. On day 14, peripheral blood samples were collected from the orbital sinus of mice from each group. After blood collection, mice were euthanized by excessive isoflurane inhalation, and the back skin and major internal organs were collected for further analysis.

### 4.13. Ethic Statement

All procedures involving BMSCs and BMMs isolation, as well as animal experiments, were approved by the Ethics Committee of the West China School of Stomatology, Sichuan University (approval number: WCHSIRB-AT-2026-435).

### 4.14. Statistical Analysis

All statistical analyses were performed using GraphPad Prism 7.0. Experiments were conducted in triplicates, or as otherwise indicated. Data are presented as the mean ± SD (standard deviation), with sample sizes (n) provided in the figure legends. Statistical comparisons between groups were made using two-sided tests, with an alpha value set at 0.05. Descriptive statistics, T-tests, one-way ANOVA, and repeated measures ANOVA were used to assess significance, with P-values < 0.05 considered statistically significant. Specific statistical tests are detailed in the figure legends.

## Data and Code Availability

## Acknowledgements

This work was supported by the National Regional Joint Key Fund project of China (No. U24A20712), the National Natural Science Foundations of China (No. 82271035, 62476183 and 82571160), and China Postdoctoral Science Foundation (No. 2025M781450). The authors thank Qiaoyu Zhou (College of Chemistry, Sichuan University) for the assistance on single crystal structure analysis, Minghai Tang (State Key Laboratory of Biotherapy, Sichuan University) for the assistance on performing the 8BA *in vivo* metabolism and pharmacokinetic determination experiments, MoDe Technology (Chengdu, China) for their assistance on MD calculations and Yiyan Testing Technology (Shanghai, China) for their assistance on HAADF-STEM.

## Author Contributions

Z.W, H.W., X.H. and H.Z. conceived this study. Z.W., H.W. and Z.H built the AI model. T.W. and Z.H. prepared the training data. C.Z. provided instructions for artificial intelligence modelling. Z.W., C.L. and T.L synthesized new nucleoside compounds and resolved their single-crystal structure. D.B. provided instructions on the evaluation of the biological properties of the materials. Z.W. and T.W. evaluated the *in vitro* and *in vivo* anti-inflammatory properties of the material and investigated the underlying molecular mechanisms.

## Conflict of Interest

The authors declare no conflict of interest.

## Extended Data

**Extended Data Fig. 1.**
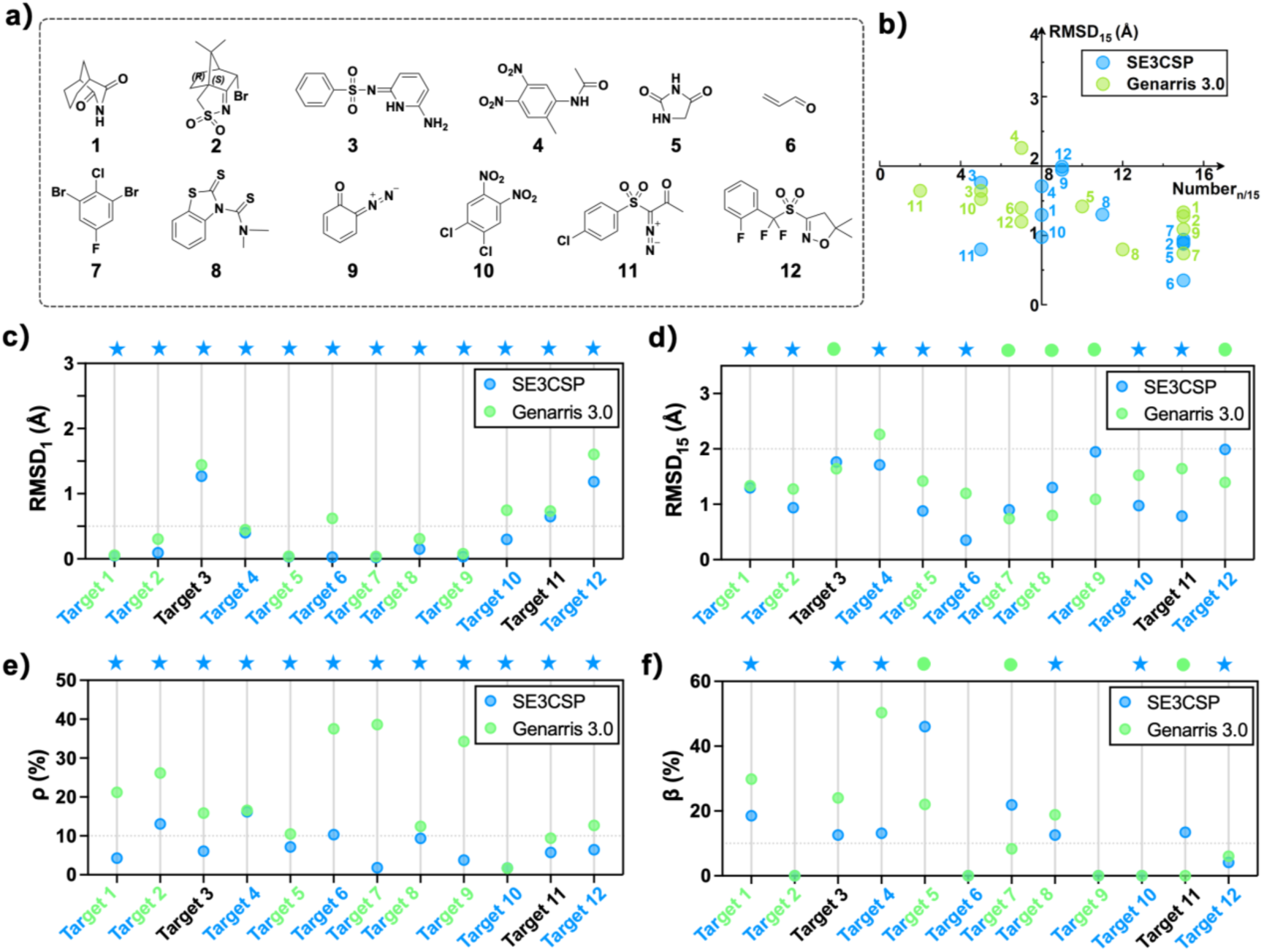
Performance comparison between SE3CSP and Genarris 3.0 for CSD blind-test candidates. a) Chemical structures of the CSD blind-test candidates. b) Two-dimensional quadrant plot of packing similarity and RMSD_15_ for structures predicted by SE3CSP and Genarris 3.0. Structures located in the lower-right quadrant represent successful predictions. c-f) Comparison of RMSD_1_, RMSD_15_, crystal density ρ and β between SE3CSP- and Genarris 3.0-predicted structures relative to the corresponding experimental structures. Blue stars indicate better performance of SE3CSP, whereas green circles indicate better performance of Genarris 3.0.

**Extended Data Fig. 2.**
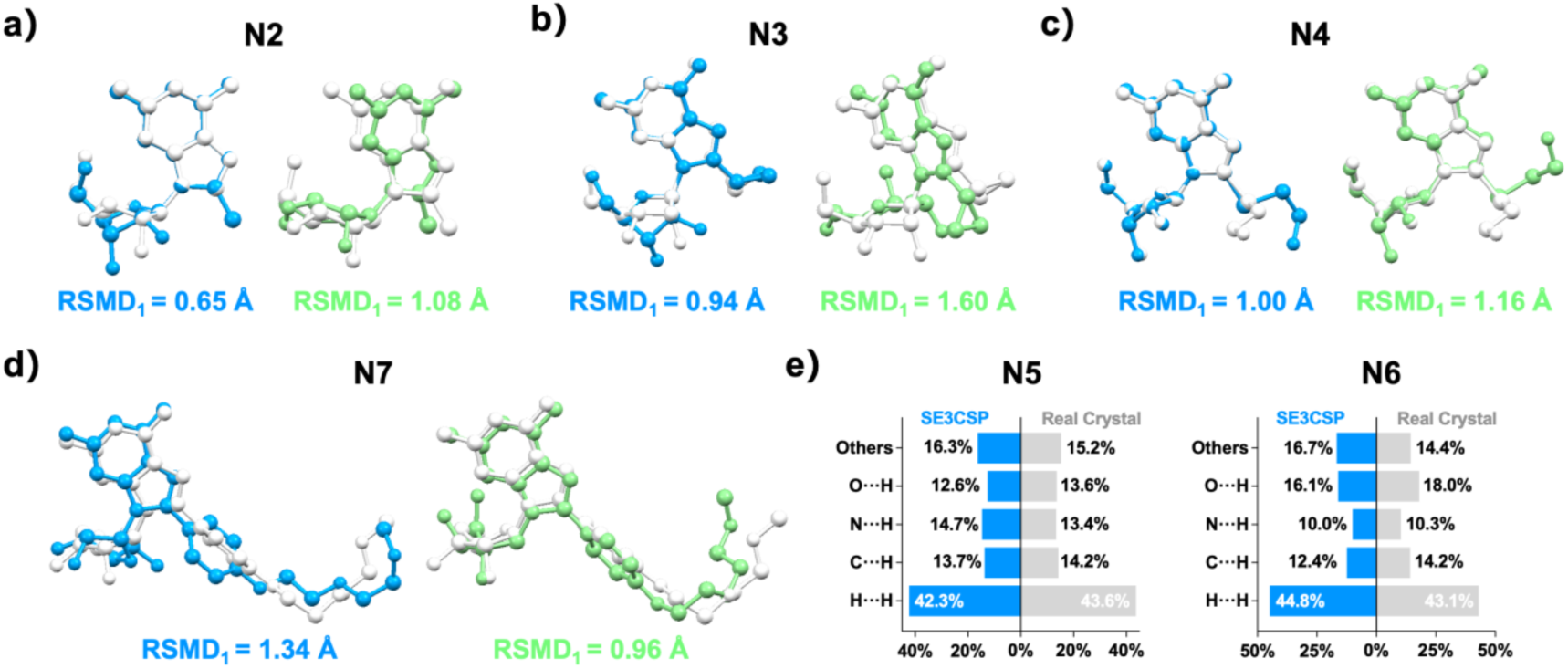
Structural comparison between SE3CSP-predicted and experimental nucleoside crystal structures. a-d) Molecular overlay of predicted and experimental crystal structures for representative cases. e) Individual atomic contact percentages obtained from Hirshfeld surface analysis for SE3CSP-predicted and experimental structures.

**Extended Data Fig. 3.**
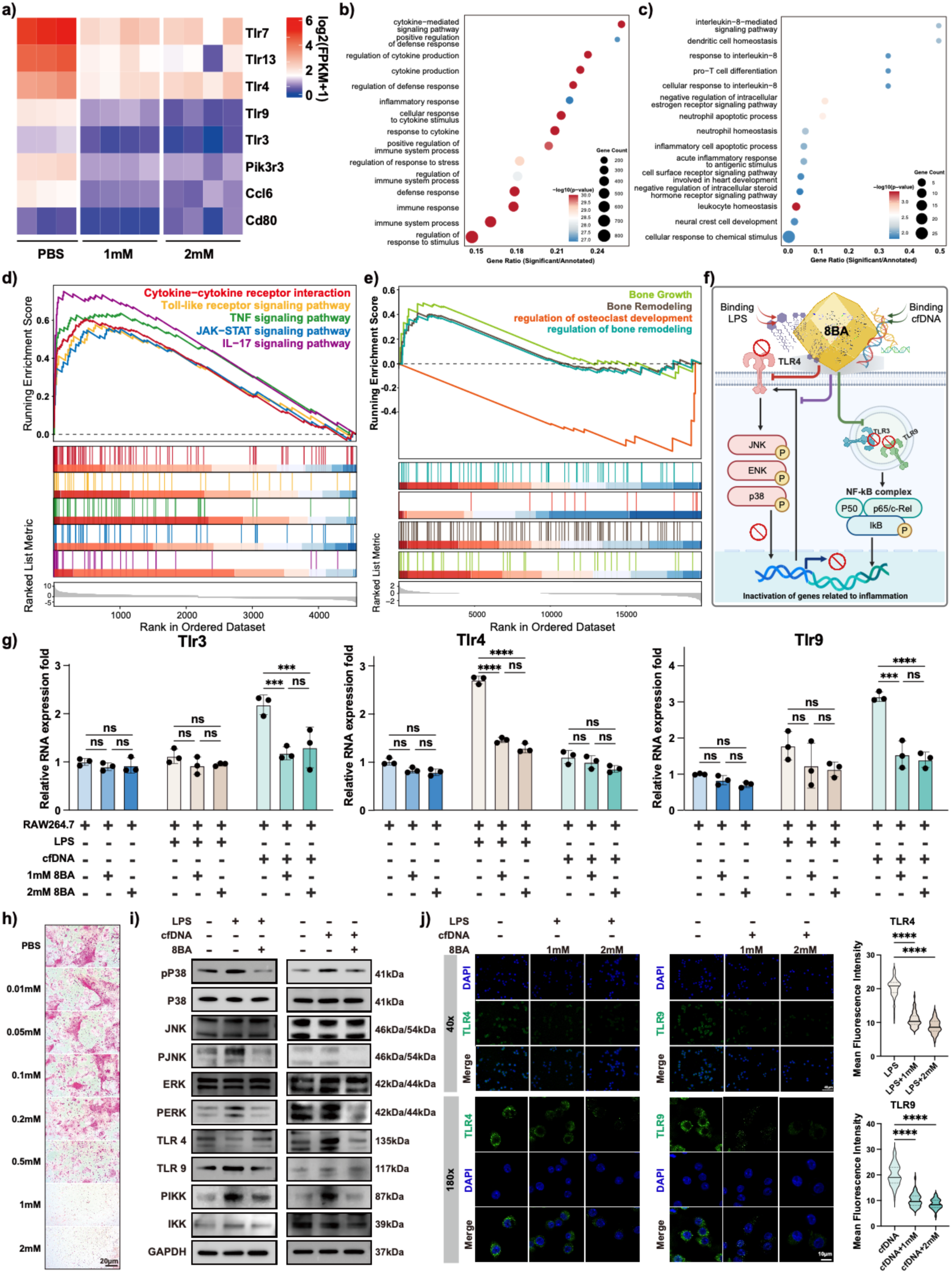
8BA suppresses TLR signaling and inhibits osteoclast differentiation. a) Heatmap showing mRNA transcription levels of TLR receptors and related molecules in PBS, 1 mM 8BA and 2 mM 8BA groups, color represents log₂(FPKM+1), n = 3. b) Bubble plot showing GO enrichment analysis of differentially expressed genes in LPS vs control. Circle size represents gene count, color represents -log₁₀(p-value). c) Bubble plot showing GO enrichment analysis of differentially expressed genes in 1 mM 8BA vs LPS groups. Circle size represents gene count, color represents -log₁₀(p-value). d) GSEA of differentially expressed genes in LPS vs control. Different colors represent different pathways. e) GSEA of differentially expressed genes in 1 mM 8BA vs LPS groups. Different colors represent different pathways. f) Schematic illustration showing that 8BA adsorbs negatively charged pathogenic molecules and simultaneously inhibits TLR receptor expression, thereby suppressing inflammation. g) Bar plot showing mRNA expression levels of Tlr3, Tlr4 and Tlr9 in RAW264.7 cells treated with LPS or cfDNA together with 1 mM or 2 mM 8BA, n = 3. h) TRAP staining of BMM cells after osteoclast induction with 0.01, 0.05, 0.1, 0.2, 0.5, 1 or 2 mM 8BA, scale bar = 20 μm. i) WB analysis of pP38, P38, JNK, pJNK, ERK, pERK, TLR4, TLR9, PIKK, IKK and GAPDH expression levels. j) Representative immunofluorescence images of TLR4 and TLR9 in RAW264.7 cells treated with LPS and 8BA at different magnifications. Right panel shows violin plot of fluorescence intensity for TLR4 and TLR9, scale bar = 40 μm (40×) or 10 μm (180×). Statistical analyses were performed using two-way ANOVA with Tukey’s multiple comparisons test in (g and j). In (g), bar graphs are shown as mean ± SD. * p < 0.05, ** p < 0.01, *** p < 0.001, **** p < 0.0001.

**Extended Data Fig. 4.**
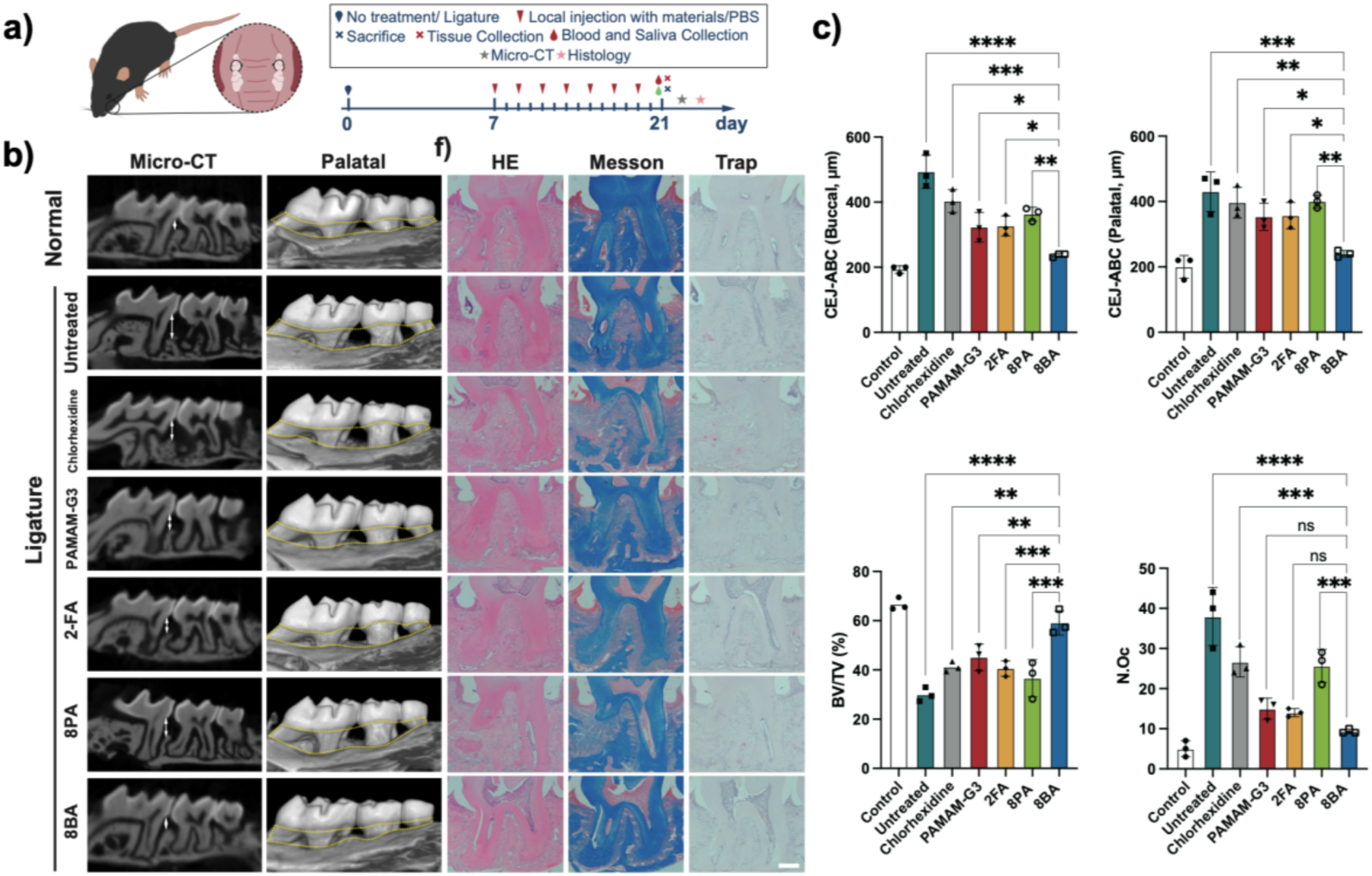
Effect of 8BA on alveolar bone regeneration in a mouse model of periodontitis. a) Schematic illustration showing the establishment of periodontitis model. b) Representative images of H&E, MESSON, TNF-α, IL-6, TLR4 and TLR9 IHC staining in tooth and periodontal tissues of non-periodontitis, periodontitis, and periodontitis groups treated with chlorhexidine, PAMAM-G3, 2FA, 8PA or 8BA. c. Bar plot showing Buccal CEJ-ABC, Palatal CEJ-ABC, BV/TV and number of osteoclast of mice in different groups, n = 3. Statistical analyses were performed using one-way ANOVA with Tukey’s multiple comparisons test in (c). In (c), bar graphs are shown as mean ± SD. * p < 0.05, ** p < 0.01, *** p < 0.001, **** p < 0.0001. Scale Bar = 200 µm.

**Extended Data Table 1.**
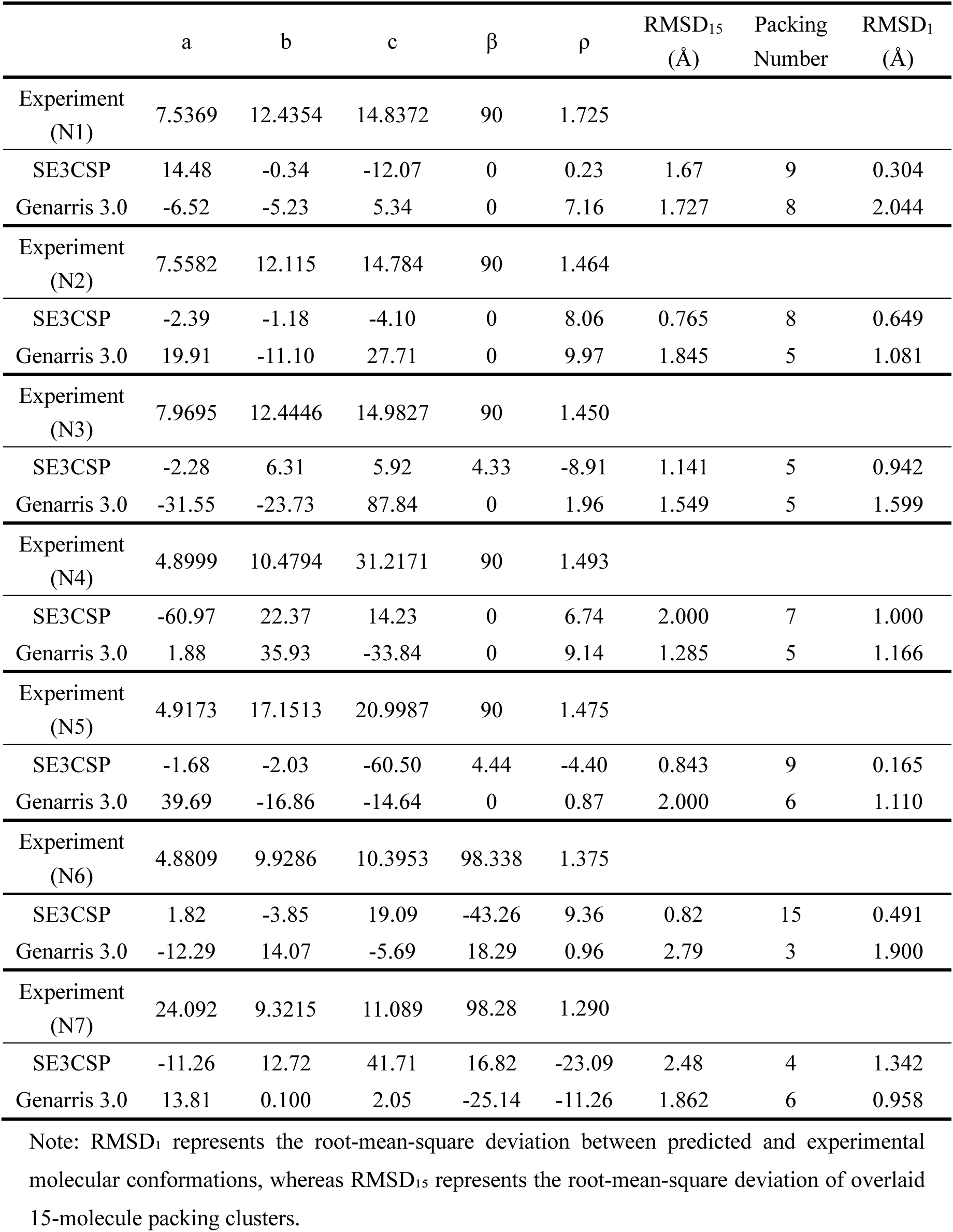
Comparison of the experimental structure and matching predictions of compounds **1-7**, in terms of the relative deviation in lattice parameters, volume and density: ((pred.−expt.)/expt.) × 100%. The RMSD_15_ and RMSD₁ are also given in Å. Experimental values for lattice parameters and density are reported in Å, °, and g/cm³, respectively.

## Notes

### Competing Interest Statement

The authors have declared no competing interest.

